# Development and calibration of the FSPM CPlantBox to represent the interactions between water and carbon fluxes in the soil-plant-atmosphere continuum

**DOI:** 10.1101/2023.04.18.537289

**Authors:** M. Giraud, S. Le Gall, M. Harings, M. Javaux, D. Leitner, F. Meunier, Y. Rothfuss, D. van Dusschoten, J. Vanderborght, H. Vereecken, G. Lobet, A. Schnepf

**Author notes:** Contributed equally to this study (M. Giraud, S. Le Gall). Supervised equally this study (G. Lobet, A. Schnepf).

## Abstract

A plant’s development is strongly linked to the water and carbon flows in the soil-plant-atmosphere continuum. Expected climate shifts will alter the water and carbon cycles and will affect plant phenotypes. Comprehensive models which simulate mechanistically and dynamically the feedback loops between a plant’s three-dimensional development and the water and carbon flows are useful tools to evaluate the sustainability of genotype-environment-management combinations which do not yet exist. In this study, we present the latest version of the open-source three-dimensional Functional-Structural Plant Model CPlantBox with PiafMunch and DuMu^x^ coupling. We simulated semi-mechanistically the development of generic C3 monocots from 10 to 25 days after sowing and undergoing an atmospheric dry spell of one week (no precipitation). We compared the results for dry spells starting on different days (day 11 or 18) and with different climates (wetter and colder against drier and warmer atmospheric and initial soil conditions). Compared with the wetter and colder climate, the dry spell with the drier and warmer climate led to a lower instantaneous water use efficiency. Moreover, the lower symplasm turgor for the drier and warmer climate limited the growth, which made the sucrose available for other processes, such as maintenance respiration. Both of these effects were stronger for the later dry spell compared with the early dry spell under the drier and warmer climate. We could thus use CPlantBox to simulate diverging emerging processes (like carbon partitioning) defining the plants’ phenotypic plasticity response to their environment.

## Introduction

Terrestrial carbon and water cycles are, amongst others, affected by plant development [16, 15, 17]. Climate change is expected to alter the global water and carbon balances [59] and thus plant phenotype and fitness [17]. A better understanding of the mechanisms driving the plant carbon- and water-fluxes, and their relations to the environment is key to selecting phenotypes and management practices, which are adapted to these altered environmental conditions so as to mitigate the negative effects of climate change and maintain or even increase food production [16, 21, 57, 17].

However, the complex spatio-temporal interactions between the water and carbon fluxes in the soil-plant-atmosphere continuum make predictions and analyses of experimental results difficult. First of all, cellular scale characteristics (like the shape and size of the xylem and phloem tissues) need to be considered to understand emerging effects seen at larger scale (like the evapotranspiration and biomass production of a field) [8, 7, 60, 13]. Moreover, feedback loops and threshold effects make the influence of plant physiological traits on the water and carbon cycles site-specific [17]. For these same reasons, we can also observe differences between short- and long-term effects of climatic shifts on the water and carbon cycles [14, 17] and experimental studies are not sufficient to predict the equilibrium of ecosystems in the distant future.

Modelling can help evaluate plant behaviour for as yet non-existing genotype-environment-management combinations and can help decipher underlying interrelated processes leading to experimental observations [58, 64, 57]. Modelling can therefore be used to investigate the benefits and drawbacks of novel crop management practices and to set objectives for plant breeders [58, 36, 57].

Three-dimensional functional structural plant models (3D FSPMs) represent emergent properties at the organism scale, which can subsequently be scaled up to the ecosystem or field scales [16, 36, 57]. FPSMs that represent the transport functions of plant tissues are especially adapted to simulate plant carbon and water fluxes in the plant and link these fluxes to [a] the water and carbon transfer in the environments (soil and atmosphere), [b] the smaller and larger scale structures of the plant, and [c] its growth and development [9, 32, 13]. For instance, axial flow (along the transport tissues) can be represented by Darcy’s law, while lateral flow (leaving or entering the transport tissues), which involves transport across selective plant cell membranes, can be represented by a membrane transport equation [39, 60, 67] and depends on pressure or osmotic potential gradients. The plant water and carbon fluxes are hence the results of [a] cellular-scale characteristics, like the number of cell layers between the vascular bundles and the epidermis [7], [b] plant-scale characteristics, like the topology of the xylem and phloem networks [32], and [c] water potentials in the soil and atmosphere and photosyntesis and local sink terms that determine sugar concentrations in the phloem. Therefore, FPSMs with transport functions should be able to predict the fluxes of carbohydrates and water from sources to sinks and determine organ growth as a function of plant-external conditions. This implies that these FSPMs should be able to reproduce phenotypic plasticity in different climates and could complement crop models in which the plant structure development, like the ratio of shoot-to-root biomass, is an empirical parameter set for a specific genotype-environment-management combination.

The quality and applicability of an FSPM depends on the processes and domains it considers: linking different mechanisms (like the flow and use of carbon), domains (soil, plant, atmosphere), and organs (root, shoot, …) via dynamic feedback loops is essential to understand the mid- and long-term effects of specific plant traits on the plant capacity to adapt to the environment [64, 36]. For instance, the short feedback loop between the plant stomatal regulation, water and carbon flows needs to be represented. Indeed, the photosynthesis (source of plant carbon) and transpiration (loss of plant water) occur through the same path and are both influenced by the stomatal opening [12, 14, 18]. This loss of water (resp. source of carbon) is interlinked with the xylem (resp. phloem) turgor pressure gradients which drive the flow in the transport tissues [60]. As the stomatal opening depends also on the leaf water status, it is itself influenced by the flow of water in the xylem [14, 18, 64]). An important longer-term feedback loop involves the plant growth: carbon and water flows regulate the partitioning of assimilated carbohydrates between the different organs (and resulting shoot-to-root ratio) [16, 32]. The growth will, in turn, indirectly influence the water and carbon fluxes in the soil-plant-atmosphere continuum by changing plant transport network topology and with this, its photosynthetic and water uptake capacities [14, 11, 21].

In this work, we present the latest implementation of CPlantBox, a 3D FSPM [67], with PiafMunch [32] and DuMu^x^ [31] coupling. The novel aspect of this CPlantBox implementation lies in the tight coupling with PiafMunch’s carbon flow module, in the implementation of CRootBox modules (like plant growth) [53] and in the addition of new modules, such as the coupled photosynthesis-transpiration-stomatal regulation and water- and carbon-limitation on growth. While the earlier version of CPlantBox could simulate the effect of a static plant, soil, and userdefined transpiration rate on the plant water and carbon fluxes, the new model can also simulate the influence of the plant water and carbon fluxes on the soil water flow, the plant growth, and the stomatal regulation. The model can consequently represent semi-mechanistically the interactions between the growth of a 3D plant (discrete structure) and the water and carbon fluxes in the soil-plant-atmosphere continuum (continuum equations). CPlantBox offers thus a holistic and precise (small space and time scale) representation of key processes (plant carbon and water uptake, flow and usage), which can be used to understand the causes of emergent behaviors (exudation rate, carbon partitioning). This new development of the model aims to be user-friendly and adapted to a wide variety of plant types.

In this paper, we present:

1. An overview of the equations implemented in the new version of CPlantBox and the schemes used to solve them.
2. An application of the model for a generic C3 monocot. The parameters were set using experimental and literature data. The model was then implemented to look at the effect on the plant’s development of a dry spell of one week of different intensities and at different plant development stages.

## Part I

### Description of the model

Figure 1 gives an overview of the different processes simulated in CPlantBox. In summary: CPlantBox aims to represent the soil-plant-atmosphere continuum and the interactions between those three domains. The environmental conditions are represented by atmospheric forcing (Figure 1.A.1) and by the soil water initial conditions. The soil water flow equations are then computed during the simulation via the the open source simulation framework DuMu^x^ [31] (Figre 1.A.3, see Section 3). The plant domain is represented by a discrete structure made of nodes linked by segments (Figure 1.A.2a, see Section 1). The atmospheric, leaf boundary layer, and soil variables affect the stomatal opening, which regulates transpiration (water sink) and photosynthesis (sucrose source) (Figure 1.A.4, see Section 2.1 and Appendix D). CPlantBox simulates the resulting xylem water flow, lateral root (and leaf) fluxes and soil-root water exchanges (Figure 1.6, see Section 2.2). CPlantBox simulates likewise the resulting mesophyll and sieve tube sucrose flow (Figure 1.5, see Section 2.3). The sucrose dynamic is also influenced by the sucrose usage (or sink), which yields the plant water- and carbon-limited growth, respiration (linked to maintenance and growth), and sucrose exudation (Figure 1.7, see Section 2.4).

**Figure 1:**
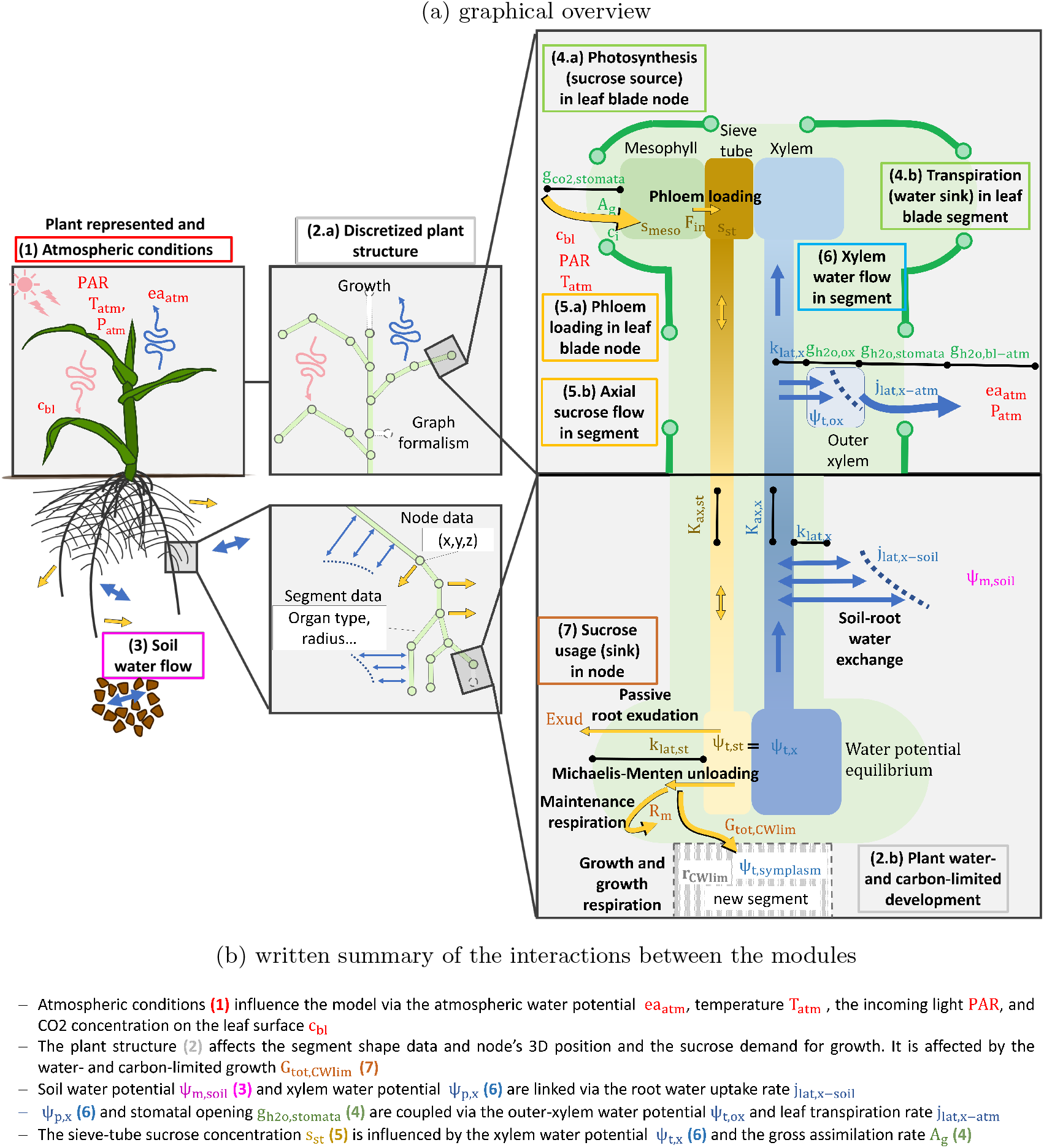
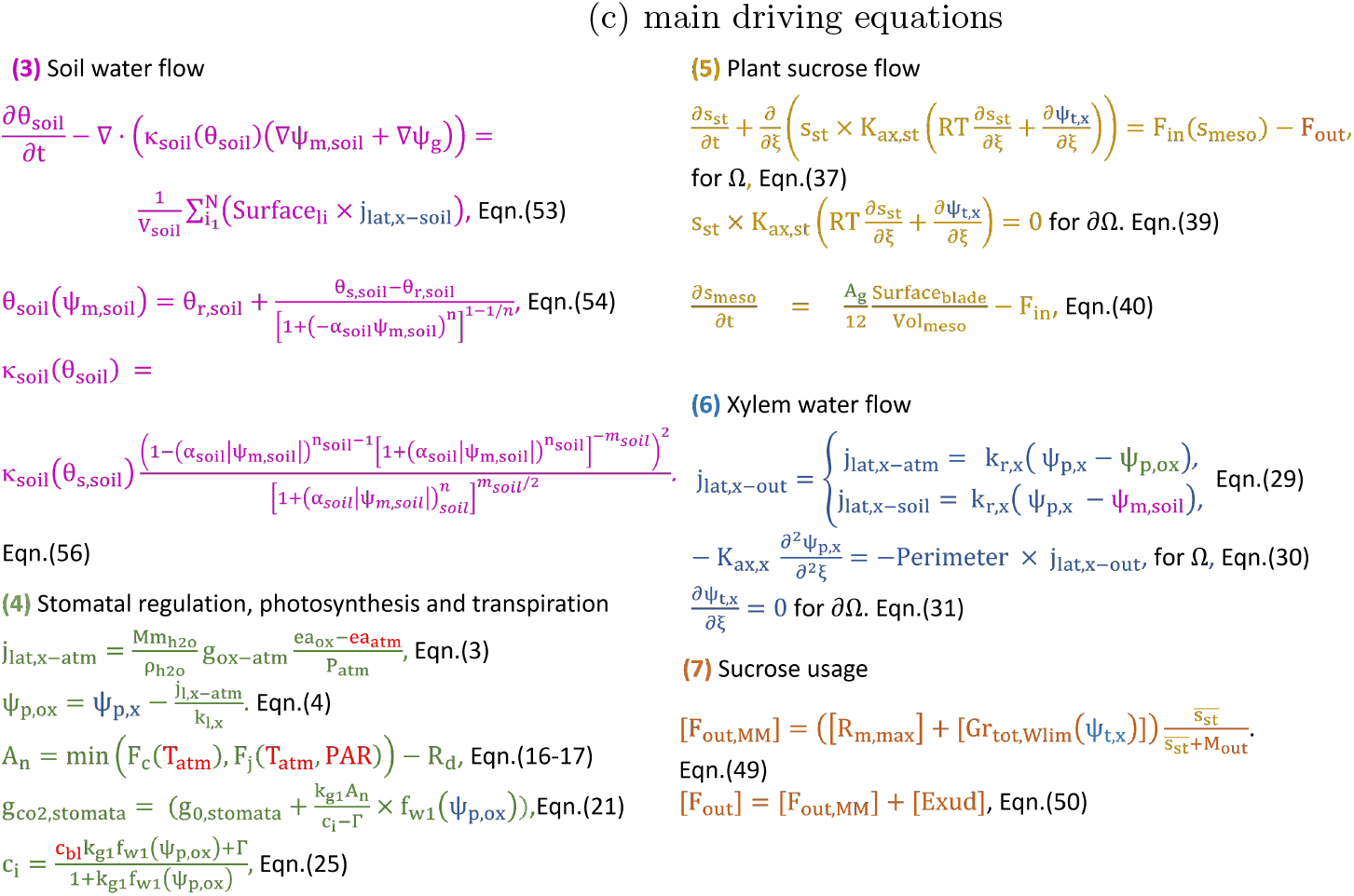
Representation of the domains and processes simulated within CPlantBox.

The main differences between the CPlantBox version presented by Zhou et al. [67] and the current version are presented in Appendix A. Briefly, we implemented a tight coupling between the phloem flow solver and CPlantBox. Then, functional modules already available for CRootBox [53] were implemented for CPlantBox; namely, the plant and soil water flow solvers. Finally, new functional modules were added; namely, the stomatal regulation-photosynthesis, sucrose usage, and water- and carbon-limited growth modules.

In the following sections, we define how plants are represented as a discrete structure in the model (Section 1). Following this, the equations of the implemented modules are listed (Sections 1-3). Finally, we describe the computation loop (Section 4). The schemes used to solve these continuum models on the discrete plant structures are presented in appendix H and a sensitivity analysis of the water-related modules and carbon-related modules are presented in appendix I. A definition of the water potential components and of the conductance or conductivity variables used in this paper as well as their notations are given in Appendix B. Tables presenting the lists of symbols used with their respective units and (when applicable) values can be found in Appendix G.2.

#### 1 Structure of the plant

As represented in Figure 2.a.1, we define a plant as an ensemble of organs. We consider the organs: “seed”, “root”, “stem”, or “leaf”. The organ class could be extended to include also other organ categories like, for instance, flowers, or fruits. The shape variables are defined with subscripts: _*seed, root, stem, leaf*_, _*st*_ (sieve tube tissues), _*x*_ (xylem tissues), _*seg*_ (segment), _*org*_ (organ), _*sheath*_ (leaf sheath), or _*blade*_ (leaf blade).

**Figure 2:**
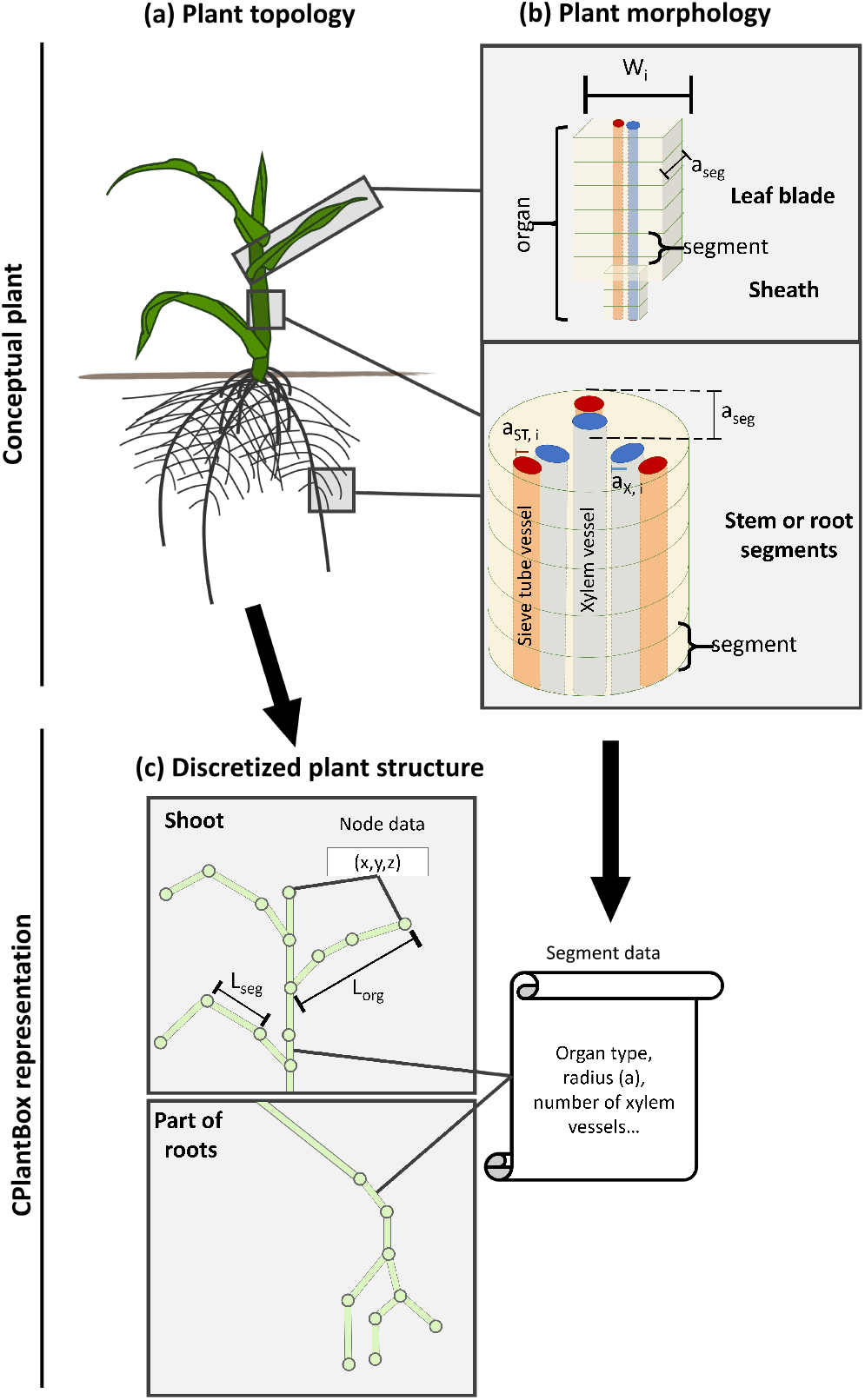
Representation of a conceptual plant’s topology (a), the morphology of its organs (b) and its discretized representation in CPlantBox (c). The plant topology is represented by the nodes’ 3-D coordinates and by their connections (segments). The morphological data are stored in arrays with one value per segment. Note: for better visibility, the cross section of the xylem and phloem vessels are not at the scale of the organs’ cross sections.

As represented in Figure 1.a.2a and Figure 2.b, plant organs are seen as a series of cylinders (root or stem segments) or cuboids (leaf segments).

As shown in Figure 2.c, the conceptual plant topology is represented in a discrete way in CPlantBox using graph formalism: the organs are made of 1-D segments, which are defined by their begin- and end-nodes (directed tree). Each node has 3-D coordinates. The orientation of the segments has an effect on the sign of the inner plant flows (see appendix H.1).

CPlantBox stores one mean value per segment of each variable defining the plant’s morphology, like cylinder radius (or cuboid thickness) *a*, and cuboid width *Wi*, both in *cm*. Although the sieve tubes and xylem tissues are not explicitly represented, their data are also stored, for instance: number of xylem vessel per cross section (*n*_*x*_) and radius (*a*_*x*_). This allows us to directly use or compute at run-time the necessary plant variables.

More information regarding the plant conceptual shape and the related equations can be found in the appendix C.

The plant growth rate can either follow an empirical function or, when the water and sucrose modules are run, the water- and sucrose-limited growth rate (*r*_*CW lim*_ in *cm d*^−1^), can be used (Figure 1.a.2b). The equations used to compute *r*_*CW lim*_ are presented in the appendix E. In brief, the initial potential (maximum) organ growth rate is user-defined. CPlantBox then computes the current potential organ growth rate, the limitation caused by the symplast turgor-pressure (*ψ*_*p,symplasm*_ in *hPa*) and by the supply of sucrose. This yields *r*_*CW lim*_, the actual growth rate. This growth is represented via the elongation of existing segments or via the creation of new segments.

The lateral growth of the xylem and phloem tissues can be represented by setting a time-dependent axial conductance, lateral conductivity, as well as a time-dependent cross sectional area of the phloem vessels (*A*_*cross,st*_ in *cm*^2^). We neglect the elastic (reversible) variation of the tissue volumes (*V ol* in *cm*^3^) according to the turgor pressures (*ψ*_*p*_ in *hPa*).

The explicit representation of the plant segment and their growth in CPlantBox is presented in more detail by Schnepf et al. [53].

#### 2 Functions within the plant

##### 2.1 Stomatal regulation, photosynthesis, and transpiration

In the following section, we present the FcVB-stomatal regulation module (see Figure 1.a.4) [20, 35, 64]. Transpiration and photosynthesis outside of the leaf blades were neglected. Outputs include [a] the net assimilation rate (*A*_*n*_ in *mmol C cm*^−2^*d*^−1^), which is the source term used by the sucrose flow module (see Eqn.(40)), and [b] the leaf outer-xylem water potential (*ψ*_*t,ox*_ in *hPa*), which is used by the xylem module (see Eqn.(29)).

###### 2.1.1 Leaf transpiration rate and stomatal hydraulic conductivity

The area-specific water vapor flow rate *j*_*lat,ox−atm*_ (*cm*^3^ *cm*^−2^ *d*^−1^) corresponds to the water flow from the outside of the vascular bundle-sheath to the point of measurement of atmospheric variables. It can be driven by the gradient of water vapor pressure (*ea* in *hPa*) or of total water potential (*ψ*_*t*_ in *hPa*). We convert *ψ*_*t*_ to *ea* using Raoult’s law [43]:

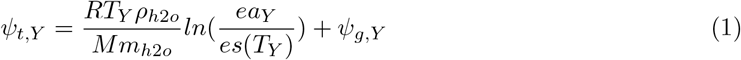

with, for the domain _*Y*_, *Mm*_*h*2*o*_ (*kg mmol*^−1^) and *ρ*_*h*2*o*_ (*kg cm*^−3^) respectively the water molar mass and density, *es*(*T*_*Y*_) (*hPa*) the saturation vapor pressure at temperature *T*_*Y*_ (*K*), the ideal gas constant *R* (*cm*^3^ *hPa K*^−1^*mmol*^−1^), and the gravitational potential *ψ*_*g,x*_ (*hPa*).

As presented in Figure 3 the water vapor has to go through several compartments, defined by their respective conductances *g*_*h*2*o,Y*_ (*mmol cm*^−2^ *d*^−1^); with _*Y*_ standing for the outer-xylem pathway to the stomata (_*ox*_), the stomata (_*stomata*_), the leaf boundary layer (_*bl*_), the canopy above the leaf (_*canopy*_), or the air between the top of the canopy and the point of measurement (_*atm*_). The total conductance between the outer xylem compartment and the point of humidity measurement (*g*_*h*2*o,ox−atm*_ in *mmol cm*^−2^ *d*^−1^) is obtained by analogy with electric resistances in series:

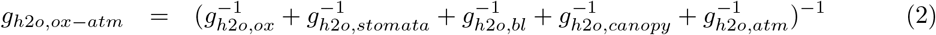

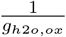 includes the resistances of different leaf elements, such as the mesophyll. The computation of *g*_*h*2*o,stomata*_ and *g*_*h*2*o,ox*_ are presented in Section 2.1.3. The evaluation of the other H_2_O conductances is presented in Appendix D.

**Figure 3:**
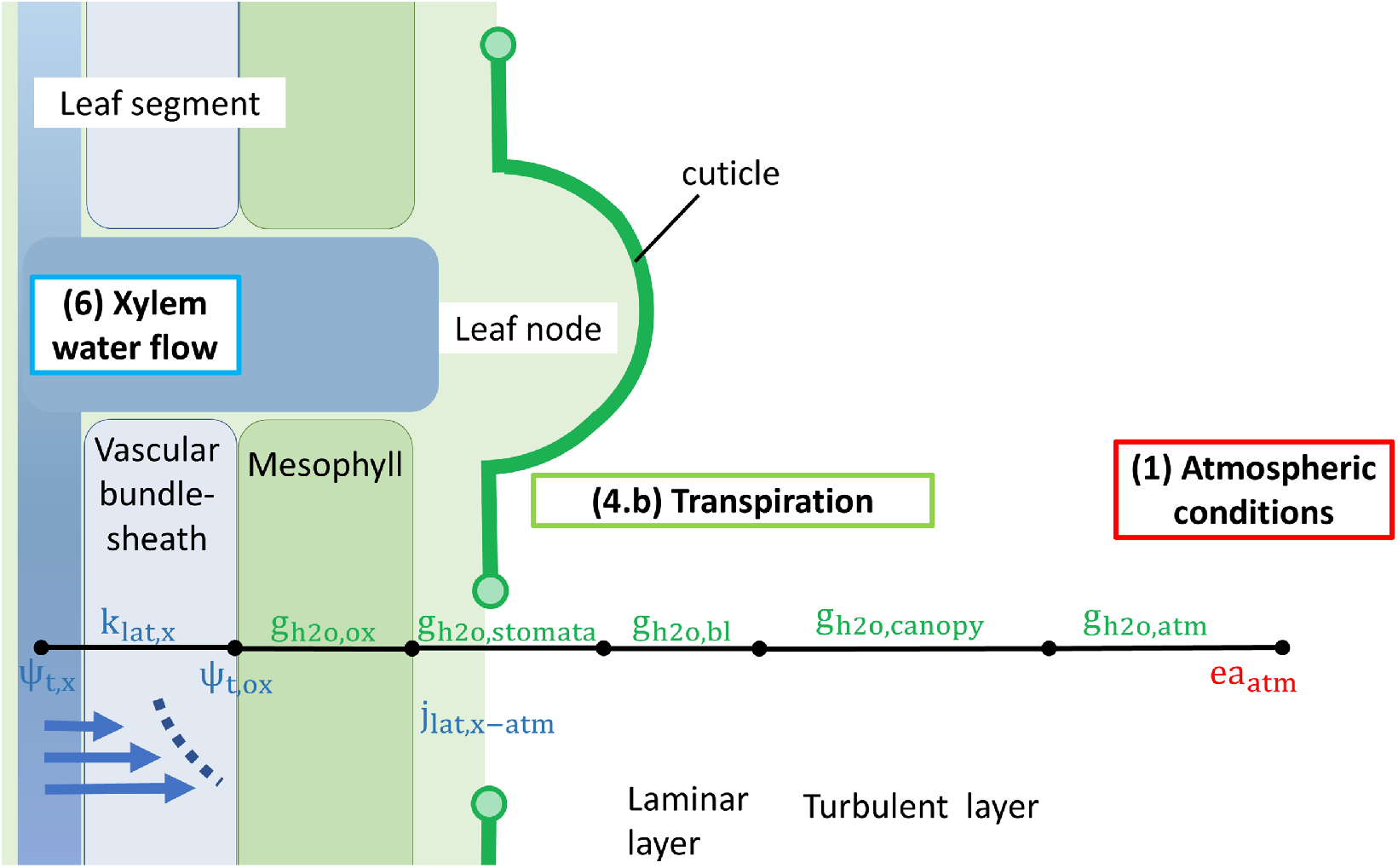
Water flow between the leaf xylem and the point of humidity measurement. The numerotation of the image sections were kept identical to those presented in Figure 1.a. The xylem total water potential (*ψ*_*t,x*_) varies along the plant segments while each segment has one mean outer-xylem water potential (*ψ*_*t,ox*_).

*j*_*lat,ox−atm*_ can then be obtained from Fick’s law applied to ideal gases [64, 44]:

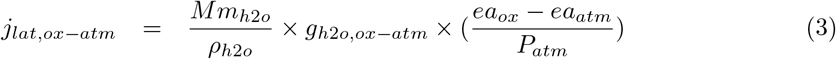

with *P*_*atm*_ (*hPa*) the atmospheric air pressure. We don’t represent the variations in water storage along the evaporation pathway. We have consequently: *j*_*lat,ox−atm*_ = *j*_*lat,x−ox*_ = *j*_*lat,x−atm*_. All the elements of the considered leaf tissue are at the same height (*ψ*_*g,ox*_ = *ψ*_*g,x*_) and we neglect the xylem osmotic potential (*ψ*_*o,x*_). *ψ*_*p,ox*_ can therefore be obtained from:

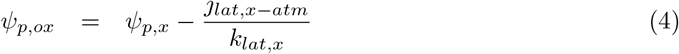

with *k*_*lat,x*_ the lateral conductivity of the vascular bundle sheath (*cm hPa*^−1^ *d*^−1^). The exact method used to compute *ψ*_*p,ox*_ is presented in Eqn.(92).

###### 2.1.2 Carbon assimilation

In this section, we present the evaluation of *A*_*n*_ as limited by the rate of carboxylation by Rubisco (*F*_*c*_ in *mmol C cm*^−2^*d*^−1^) and by the rate of RuBP regeneration via electron transport (*F*_*j*_ in *mmol C cm*^−2^*d*^−1^).

###### 2.1.2.1 Rubisco activity at the sites of carboxylation

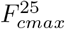 (*mmol cm*^−2^ *d*^−1^) corresponds to the maximum *F*_*c*_ for *T* = 298.15 *K*. Qian et al. [47] defined 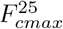 according to the leaf chlorophyll content (*Chl* in *mmol cm*^−2^):

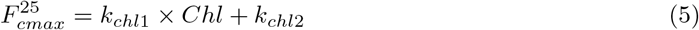

with *k*_*chl*1_ and *k*_*chl*2_ fitting parameters representing the empirical relationship between 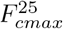 and *Chl. k*_*chl*1_ and *k*_*chl*2_ can be set according to the assimilation rate when *F*_*c*_ is limiting for photosynthesis. *Chl* is currently an input parameter and can be used to represent the effect of plant nitrogen uptake.

*F*_*cmax*_ (*mmol cm*^−2^ *d*^−1^) is the maximal *F*_*c*_ (*mmol cm*^−2^ *d*^−1^) at a temperature *T* :

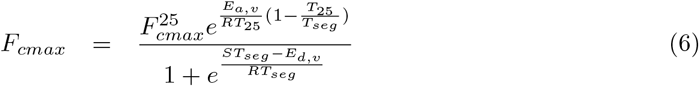

*E*_*a,v*_ and *E*_*d,v*_ (*mJ mmol*^−1^) are respectively the activation and deactivation energy for *F*_*cmax*_. *S* (*mJ mmol*^−1^ *K*^−1^) is an entropy term.

*F*_*c*_ is affected by G*\** (*mmol CO*_2_ *mmol air*^−1^), the CO_2_ compensation point (equilibrium CO_2_ level reached when *A*_*n*_ = 0) in the absence of mitochondrial respiration (*R*_*d*_ in *mmol cm*^−2^ *d*^−1^). G*\** is obtained from Eqn.(7).

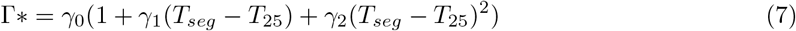

*γ*_0_ (*mmol CO*_2_ *mmol air*^−1^), *γ*_1_ (*K*^−1^), and *γ*_2_ (*K*^−2^) are empirical coefficients which define the temperature dependence of G*\**.

*F*_*c*_ follows a Michaelis-Menten function and depends on the CO_2_ and O_2_ molar fractions in the substomatal cavity, respectively *c*_*i*_ (*mmol CO*_2_ *mmol air*^−1^) and *o*_*i*_ (*mmol O*_2_ *mmol air*^−1^). It depends likewise on the Michaelis coefficients for CO_2_ and O_2_, respectively *M*_*co*2_ and *M*_*o*2_ (*mmol mmol*^−1^). *M*_*co*2_, *M*_*o*2_ (and *R*_*d*_) can be calculated with the following equation:

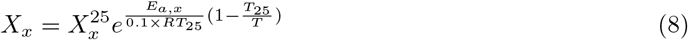

where *E*_*a,x*_ is the activation energy for *X*_*x*_ (either *M*_*co*2_, *M*_*o*2_, or *R*_*d*_) in *mJ mmol*^−1^, and *X*^25^ is the value of *X*_*x*_ for *T* = 298.15 *K*. The factor 0.1 is used to go from *hPa cm*^3^ *K*^−1^ *mmol*^−1^ to *mJ K*^−1^ *mmol*^−1^. When *A*_*g*_ *< R*_*d*_, we have a negative sucrose source in the mesophyll (see Eqn.(40)). In such cases, *R*_*d*_ is also limited by the amount of sucrose available in the mesophyll and can be lower than the value obtained from Eqn.(8).

*F*_*c*_ can then be computed using Eqn.(9):

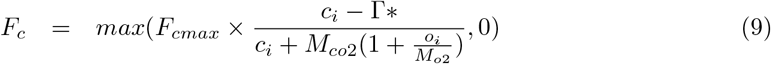

The computation of *c*_*i*_ is presented in Section 2.1.4.

###### 2.1.2.2 Electron transport rate

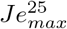 (*mmol cm*^−2^ *d*^−1^) is the the maximum electron transport rate for *T* = 298.15 *K*. It is assumed to be proportional to 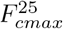 [64]:

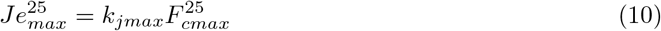

with *k*_*jmax*_ (−) a set coefficient which can be defined according to the light saturation point for photosynthesis under well-watered conditions. *Je*_*max*_ (*mmol cm*^−2^ *d*^−1^) is the the maximum electron transport rate and is computed with the following equation:

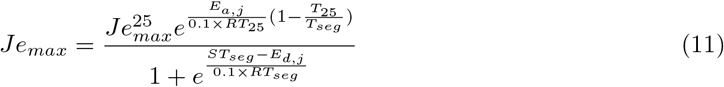

with *E*_*a,j*_ and *E*_*d,j*_ (*mJ mmol*^−1^), respectively the activation and deactivation energy for *Je*_*max*_. We can then obtain *Je* (*mmol cm*^−2^ *d*^−1^), the electron transport rate for a value of photosynthetically active radiation absorbed (*PAR* in *mmol photons cm*^−2^ *d*^−1^):

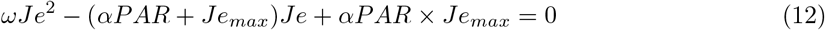

with *ω* (−) a parameter that determines the shape of the parabola and *α* (−) the quantum yield of whole-chain electron transport.

*Je* corresponds to the smaller root of the quadratic equation [5]:

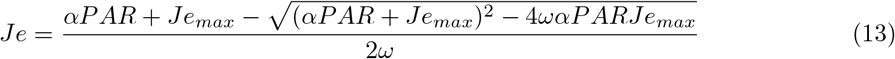

From *Je*, we can compute *F*_*j*_ :

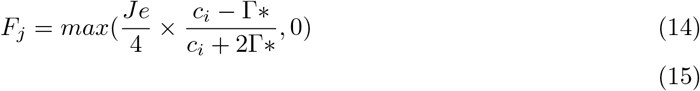

###### 2.1.2.3 Gross and net assimilation rates

The gross and net CO_2_ assimilation rates (respectively *A*_*g*_ and *A*_*n*_ in *mmol cm*^−2^ *d*^−1^) correspond to the quantity of sucrose gained by the plant per unit of leaf exchange surface and time. They are computed from *F*_*c*_ (Eqn.(9)), *F*_*j*_ (Eqn.(14)), and *R*_*d*_ (Eqn.(8)):

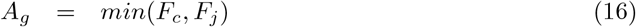

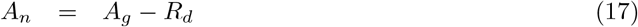

*A*_*n*_ is then used as source term for the sucrose flow (see Section 2.3.2).

###### 2.1.3. Leaf molar conductance

The molar conductances of the domaine _*Y*_ to CO_2_ and H_2_O (respectively *g*_*co*2,*Y*_ and *g*_*h*2*o,Y*_ in *mmol cm*^−2^*d*^−1^) correspond to the flow of gaseous CO_2_ (resp. H_2_O) per unit of CO_2_ concentration (resp. water vapor pressure). *g*_*co*2,*stomata*_ depends on the gradient between *c*_*i*_ and the CO_2_ compensation point (Γ in *mmol CO*_2_ *mmol air*^−1^). Γ is obtained from Eqn.(18).

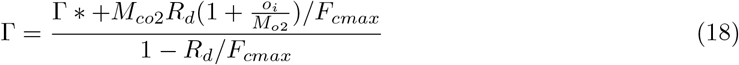

A low *ψ*_*t,ox*_ will lead to lower *g*_*co*2,*stomata*_ and *g*_*h*2*o,ox*_. This response is represented empirically via a dimensionless water scarcity factor (*f*_*w*1_):

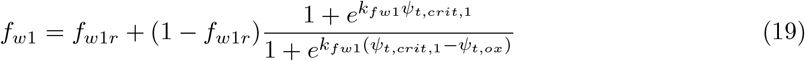

with *ψ*_*t,crit*,1_ (*hPa*) the critical water potential. *k*_*fw*1_ (−) is a sensitivity parameter. A higher value of *k*_*fw*1_ leads to a stronger variation of *f*_*w*1_ around *ψ*_*t,crit*,1_, whereas a lower value leads to a more gradual response.

*g*_*h*2*o,ox*_ (*mmol cm*^−2^*d*^−1^), the conductivity to water of the pathway between the xylem membrane and the substomatal cavity, is computed from its maximum value (*g*_*h*2*o,ox,max*_, assumed constant) and the water scarcity factor *f*_*w*1_: *g*_*h*2*o,ox*_ = *g*_*h*2*o,ox,max*_ *× f*_*w*1_.

*g*_*h*2*o,stomata*_ and *g*_*co*2,*stomata*_ are computed following the model of Tuzet et al. [64]:

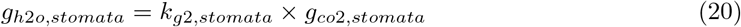

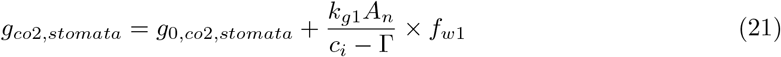

with *g*_0,*co*2,*stomata*_ (*mmol cm*^−2^*d*^−1^) the residual stomatal opening. *k*_*g*2_ accounts for the H_2_O to CO_2_ ratio of molecular diffusivity—*k*_*g*2,*stomata*_ = 1.6 [64]. As opposed to the model of Leuning [35], the model of Tuzet et al. [64] model uses *c*_*i*_ and not *c*_*bl*_ (*mmol CO*_2_ *mmol air*^−1^), the CO_2_ molar fraction at the leaf surface. Indeed, *c*_*i*_ was shown to have a stronger influence on *g*_*co*2,*stomata*_ [14]. *k*_*g*1_ (−) is a fitting parameter representing the effect of *A*_*g*_ on *g*_*co*2,*stomata*_. It can be computed from the expected 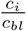ratio when *f*_*w*1_ = 1 using Eqn.(25) presented in the next section.

###### 2.1.4 Substomatal CO_2_ molar fraction

Following an Ohm’s law adaptation of Fick’s law of diffusion, *c*_*i*_ (*mmol mmol*^−1^) can be calculated with the “CO_2_ supply function” [14, 18]:

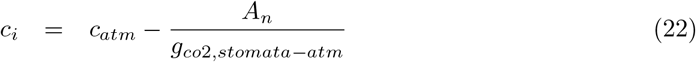

with *c*_*atm*_ (*mmol mmol*^−1^) the CO_2_ molar fraction at the point of measurement. However, when *A*_*g*_ is low, our fixed-point iteration solving scheme (see Section 4) does not converge and gives unrealistic *c*_*i*_ value (*c*_*i*_ *≤* 0 or *c*_*i*_ *≫ c*_*s*_). For this reason, similarly to Dewar [18], we use the CO_2_ molar fraction on the leaf surface (*c*_*bl*_ in *mmol mmol*^−1^).

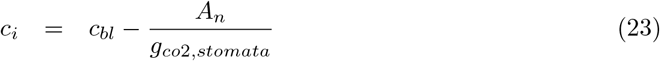

We then combine Eqn.(21) and Eqn.(23) and we neglect *g*_0,*co*2,*stomata*_:

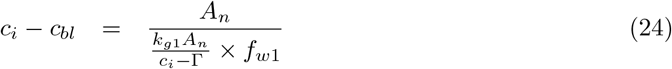

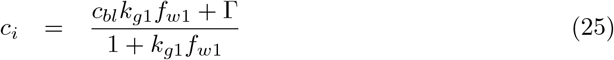

If we assume that *c*_*bl*_ and *k*_*g*1_ are set parameters and G *≪ c*_*bl*_, the *c*_*i*_ value then mainly depends on *f*_*w*1_ (−), the water scarcity factor of the stomatal model. This follows field observations, where 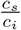 is usually seen to be constant, except in case of water scarcity [14].

##### 2.2 Xylem water flow

Darcy’s law is seen as a good approximation to compute the axial flux of water in the xylem (*J*_*ax,x*_ in *cm*^3^ *d*^−1^) [50]. Moreover, we assume that the xylem solution has physical characteristics similar to that of pure water. We therefore neglect *ψ*_*o,x*_ (*hPa*), the xylem osmotic potential. We thus have *ψ*_*t,x*_ = *ψ*_*p,x*_ + *ψ*_*g,x*_, with *ψ*_*t,x*_ the total xylem solution potential, *ψ*_*p,x*_ the xylem turgor pressure and *ψ*_*g,x*_ the gravitational potential, all in *hPa*. For a straight xylem segment we obtain:

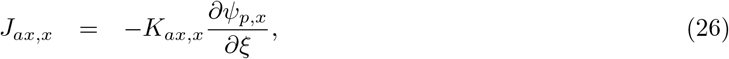

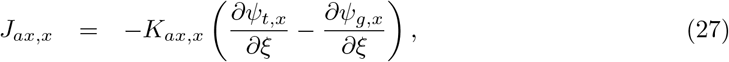

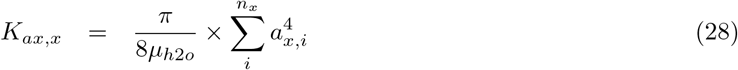

*K*_*ax,x*_ (*cm*^4^ *hPa*^−1^*d*^−1^) is the xylem intrinsic axial conductance and *ξ* (*cm*) corresponds to the local axial coordinate of the plant. *µ*_*h*2*o*_ (*hPa d*) is the temperature-dependent viscosity of pure water. *a*_*x,i*_ (*cm*) is the radius of the xylem vessel number *i* in the segment cross-section, *n*_*x*_ is the number of xylem vessels in the cross-section.

The sink and source for water are given by the lateral water flow (*j*_*lat,x−out*_ in *cm d*^−1^), which corresponds to the xylem-soil (*j*_*lat,x−soil*_) or xylem-atmosphere (*j*_*lat,x−atm*_) water exchange:

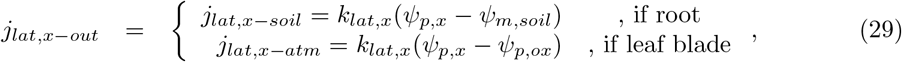

where *k*_*lat,x*_ (*cm hPa*^−1^ *d*^−1^) is the lateral hydraulic conductivity of the root cortex (for roots) or of the vascular bundle-sheath (for leaves). *ψ*_*m,soil*_ is the soil matric water potential in *hPa. ψ*_*m,soil*_ is given by the soil water flow module (see Section 3) and *ψ*_*p,ox*_ is given by the FcVB-stomatal regulation module (see Section 2.1, Eqn.4). CPlantBox offers several options for setting *k*_*lat,x*_ and *K*_*ax,x*_ (constant, age dependent, organ type dependent…). We do not currently represent the effect of cavitation on the xylem conductivity. *j*_*lat,x*−out_ > 0 represent a water loss for the plant.

We neglect the variation of water storage in the plant. Mass balance gives us thus the following driving equation for the xylem solution flow (see Figure 1.a.6):

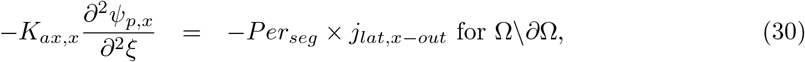

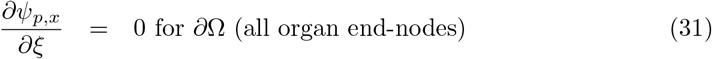

with *Per*_*seg*_ (*cm*) the perimeter of the plant exchange zone, Ω the domain considered (here the plant), and *∂*ζ, the domain’s boundary (here: the tip or end-node of each organ). Thus, the soil (Section 3, Figure 1.a.3), xylem (Figure 1.a.6), and photosynthesis-stomatal opening subproblems (Section 2.1.1, Figure 1.a.4b) are coupled via Eqn.(30).

##### 2.3 Sucrose balance

In the following section, we present the equations defining the plant sucrose dynamic (Figure 1.a.5), yielding the sucrose concentration in the sieve tube (*s*_*st*_ in *mmol sucrose cm*^−3^) and in the leaf mesophyll (*s*_*meso*_ in *mmol sucrose cm*^−3^). The flow and concentration of sucrose in other tissues (such as the parenchyma) is not explicitly represented. The module also yields [*F*_*out*_] (*mmol sucrose cm*^−3^*d*^−1^), the *s*_*st*_ loss rate, which gives the exudation rate and the plant water- and carbon-limited growth (see Section 2.4).

###### 2.3.1 Sucrose in the phloem

Similarly to Section 2.2, we compute *J*_*ax,st*_ (*cm*^3^ *d*^−1^), the sieve tube axial solution flow, by implementing Darcy’s law (see Eqn.(26)). We therefore obtain [60]:

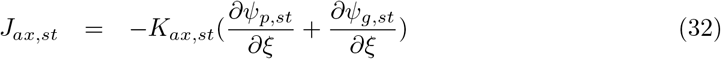

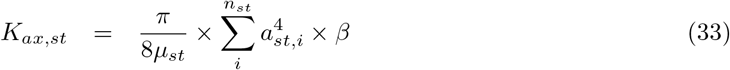

*K*_*ax,st*_ (*cm*^4^ *hPa*^−1^*d*^−1^) is the sieve tube axial intrinsic conductance and *ψ*_*p,st*_ (*hPa*) is the sieve tube hydrostatic pressure.

*µ*_*st*_ (*hPa d*) is the viscosity of the sieve tube solution and it is computed from the temperature and sucrose concentration according to the method of Mathlouthi and G’enotelle [38, Eqn. (6.29)] as implemented by Lacointe and Minchin [32]. *a*_*st,i*_ (*cm*) is the radius of the sieve tube number *i* in the segment cross-section, *n*_*st*_ is the number of sieve tubes in the cross-section.

Following the recommendation of Thompson and Holbrook [60], we added the parameter *β* (−) to the computation of *K*_*ax,st*_ to represent the ratio of axial conductance with sieve plates to that without. *β* can be evaluated via empirical data or from images of phloem and sieve plates [60].

This parameter is not needed for the xylem tissues as the opening in its cells’ end walls are much larger [42]. CPlantBox offers several options for defining *a*_*st*_ (constant, organ type dependent…).

We assumed water equilibrium between the xylem and phloem (see appendix F for more explanations). We therefore obtain:

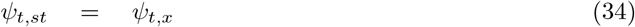

We assume that the phloem has a semi-permeable membrane (permeable to water but not to sucrose). The total potential in the phloem, *ψ*_*t,st*_, is equal to the sum of sieve tube solution’s hydrostatic (*ψ*_*p,st*_, *hPa*), gravitational (*ψ*_*g,st*_, *hPa*), and osmotic potential (*ψ*_*o,st*_, *hPa*). We consequently obtain:

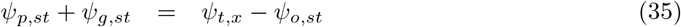

We consider that sucrose makes up the largest fraction of solutes in the sieve tube solution [15]. The *ψ*_*o,st*_ can therefore be computed according to van’t Hoff relation: *ψ*_*o,st*_ = *−RT*_*seg*_*s*_*st*_. With *T*_*seg*_ the segment’s temperature in *K*. As explained by Hall and Minchin [24] and Cabrita [8, Section 2.2.3.1], the van’t Hoff relation may not be appropriate when *s*_*st*_ > 0.5 *mmol sucrose cm*^−3^. We kept nonetheless that definition of *ψ*_*o,st*_ for this first implementation of the model.

Eqn.(32) becomes:

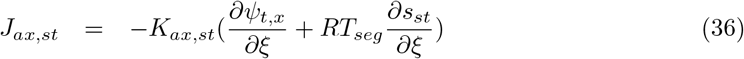

Using the relations defined above, we do not need so solve another equation for *ψ*_*p,st*_ but can at all times compute it from other known state variables, namely *ψ*_*t,x*_ (see Eqn.(30)) and *s*_*st*_.

Assuming then that sucrose is transported within the phloem by advection only, with no lateral gradient, constant initial conditions and no-flux boundary conditions, we obtain:

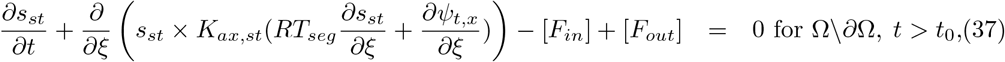

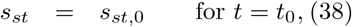

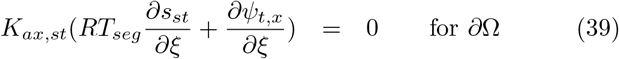

with [*F*_*in*_] and [*F*_*out*_] (*mmol sucrose cm*^−3^*d*^−1^), respectively the source and sink of *s*_*st*_ (see sections 2.3.2 and 2.4 respectively), and *t*_0_ the time at the beginning of the simulation.

###### 2.3.2 Sucrose in the mesophyll

Like *s*_*st*_, we solve *s*_*meso*_ (*mmol sucrose cm*^−3^), the sucrose concentration in the leaf source compartments (considered to be the mesophyll cells). As represented in Figure 1.a.5a, *A*_*n*_ is obtained from the photosynthesis module (see Section 2.1) and leads to a variation of *s*_*meso*_. Then occurs an active phloem loading from the mesophyll to the sieve tube ([*F*_*in*_] in *mmol sucrose cm*^−3^ *d*^−1^) [15], defined according to Stanfield and Bartlett [58]. We do not simulate axial flow within the mesophyll compartments. In summary:

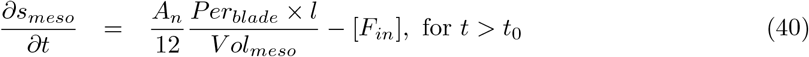

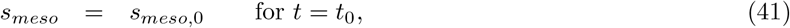

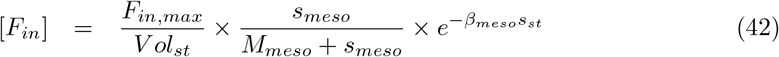

*Per*_*blade*_ (*cm*) and *V ol*_*meso*_ (*cm*^3^) are respectively the leaf blade perimeter and mesophyll volume. The factor 12 allows us to go from *mmol C d*^−1^ to *mmol sucrose d*^−1^. *F*_*in,max*_ (*mmol sucrose d*^−1^) is the maximum loading rate into the sieve tube, *β*_*meso*_ (−) is a factor indicating the strength of the *s*_*st*_ down-regulation for loading. *M*_*meso*_ (*mmol sucrose cm*^−3^) is the Michaelis-Menten coefficient for the sucrose loading. When *A*_*n*_ *<* 0, the sucrose in the mesophyll is used for mitochondrial respiration (*R*_*d*_, see Eqn.(8)).

##### 2.4 Sucrose usage

In this section, we present how to compute the maximal or water-limited sucrose usage rates for each sink categories. From this we obtain the total sucrose sink term ([*F*_*out*_] in *mmol sucrose cm*^−3^ *d*^−1^) used in the *s*_*st*_ balance equation (see Eqn.(37) and Figure 1.a.7). We also present how [*F*_*out*_] is used to compute the realized (*s*_*st*_-limited) sinks (Figure 1.a.2b).

In the plants, *s*_*st*_ can be used for growth (*Gr*, in *mmol sucrose d*^−1^), root exudation (*Exud*, in *mmol sucrose d*^−1^) and respiration [19, 62, 21]. *s*_*st*_ respiration can be divided in two conceptual categories: growth respiration (*R*_*gr*_, in *mmol sucrose d*^−1^), which is linked to *Gr*; and maintenance (or residual) respiration (*R*_*m*_, in *mmol sucrose d*^−1^). This conceptual representation follows the growth-maintenance paradigm [51, 62, 2]. Other sucrose sinks, notably starch synthesis, are not represented (discussed in Section 3). Each carbon sink *X* (*mmol sucrose d*^−1^) can also be given in concentration units (with [*X*] in *mmol sucrose cm*^−3^ *d*^−1^), obtained from 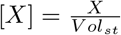, with *V ol*_*st*_ (*cm*^3^) the volume of the sieve tube.

###### 2.4.1 Exudation

The root exudation rate (*Exud*, in *mmol sucrose d*^−1^) is assumed to be a passive process [21], dependent on the lateral sucrose gradient:

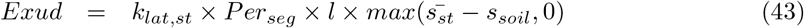

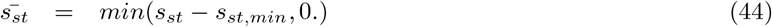

with *k*_*lat,st*_ (*cm d*^−1^) the lateral sieve tube conductivity for sucrose. Similarly to *k*_*lat,x*_, CPlantBox offers several options for setting *k*_*lat,st*_ (constant, root order dependent…). *s*_*soil*_ (*mmol sucrose cm*^−3^) is the mean soil carbon concentration in equivalent sucrose. *l* (in *cm*) is the evaluated length. *s*_*st,min*_ (*mmol sucrose cm*^−3^) is the minimum sucrose concentration below which no usage of sucrose occurs. This parameter allows the user to define more easily a minimum sucrose concentration to follow experimental observations and to respect the necessary conditions for water equilibrium between xylem and phloem (see appendix F).

###### 2.4.2 Water-limited growth and growth respiration

The water-limited (or potential) growth-related sink (*G*_*tot,W lim*_, in *mmol sucrose d*^−1^) corresponds to the potential carbon used for growth (*Gr*_*W lim*_) and growth related respiration (*R*_*gr,W lim*_). It is computed thus:

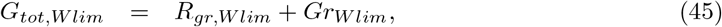

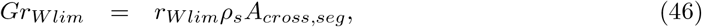

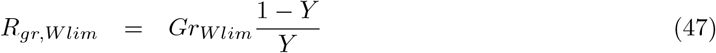

with *Y* (−) the sucrose use efficiency for growth. *ρ*_*s*_ (*mmol sucrose cm*^−3^) is the volumetric structural carbon content of this tissue in unit of sucrose. *A*_*cross,seg*_ (*cm*^2^) is the cross-sectional area of the segment. *r*_*W lim*_ (*cm d*^−1^) is the water limited growth rate, computed via Eqn.(68).

###### 2.4.3 Maximal maintenance respiration

*R*_*m,max*_ (*mmol sucrose cm*^−3^ *d*^−1^) corresponds to the maximal (non-carbon limited) value of *R*_*m*_. It represents for instance the sucrose usage to replace the structural carbon (re-synthesis), the phloem leakage and futile cycles [62, 32]. *R*_*m,max*_ is obtained empirically from Eqn.(48):

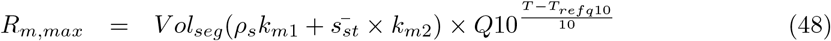

with *V ol*_*seg*_ (*cm*^3^) the volume of the plant tissues. *k*_*m*1_, *k*_*m*2_ are fitting parameters defining *R*_*m,max*_ according to, respectively, the structural (for wall re-synthesis) and non-structural carbon content. Following the example of Gauthier et al. [22], we added the short-term effect of temperature on *R*_*m,max*_ via *Q*10 (−) and *T*_*refq*10_ (*K*) [2]. *Q*10 is a coefficient which is widely used in the scientific literature and which defines the temperature sensitivity of chemical reactions: higher *Q*10 value leads to steeper increase with temperature. *T*_*refq*10_ defines the reference temperature: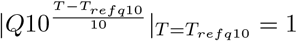. As other modules are added to CPlantBox (like nitrogen uptake and flow) their related respiration may be added to the computation of *R*_*m,max*_.

###### 2.4.4 Sucrose-limited sinks

[*F*_*out,MM*_] (*mmol sucrose cm*^−3^ *d*^−1^) is the sucrose usage rate for growth and maintenance. It follows a saturation-dynamic and is computed from the Michaelis-Menten respons fonction [22, 28, 32]:

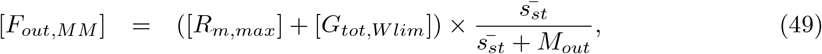

with *M*_*out*_ the Michaelis-Menten coefficient. After adding the passive *Exud*, we obtain [*F*_*out*_] (*mmol sucrose cm*^−3^ *d*^−1^) the total sucrose usage rate:

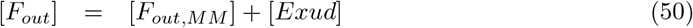

A high increase in *s*_*st*_ will only affect the maintenance and growth rate if the sink cells are not saturated, while the exudation will not be demand-limited.

*F*_*out,MM*_ gives us the value of [*R*_*m*_] and [*G*_*tot,CW lim*_] (*mmol sucrose cm*^−3^ *d*^−1^), the realized sucrose usage rate for maintenance and growth respectively :

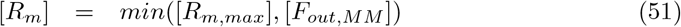

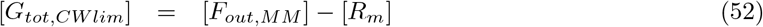

Eqn. (51) indicates that maintenance has priority over growth, which is a common assumption [51, Fig. 1].

#### 3 Soil water flow

The variation in soil water content 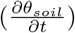 is computed thanks to the DuMu^x^ module (see Figure 1.a.3) using the Richards equation, which describes the movement of water in an unsaturated soil [49, 31]:

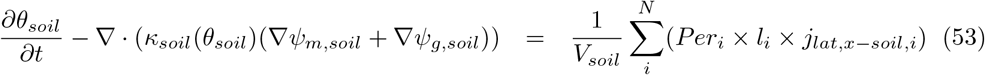

with 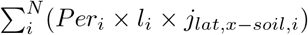, the root-soil water exchange for all roots _*i*_ in the considered soil domain (Neumann boundary condition obtained from Section 2.2), *V*_*soil*_ (*cm*^3^) the volume of the considered soil domain, and *κ*_*soil*_(*θ*_*soil*_) (in *cm*^2^ *hPa*^−1^ *d*^−1^) the soil conductivity for a specific water content.

The soil water content (*θ*_*soil*_ in *cm*^3^ *cm*^−3^) and *κ*_*soil*_(*θ*_*soil*_) are obtained from the van Genuchten-Mualem equations [41, 66]:

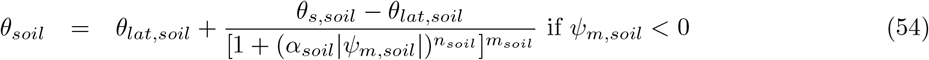

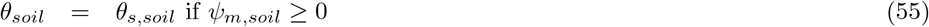

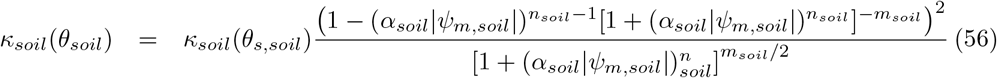

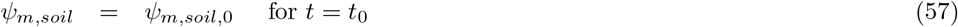

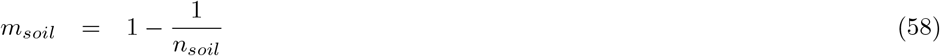

with *α*_*soil*_ (*hPa*^−1^) a parameter related to the inverse of the air-entry suction, *n*_*soil*_ and *m*_*soil*_ (−) parameters representing the soil pore-size distribution, *θ*_*s,soil*_ and *θ*_*lat,soil*_ (*cm*^3^ *cm*^−3^) the saturated and residual soil water content, *κ*_*soil*_(*θ*_*s,soil*_) (in *cm*^2^ *hPa*^−1^ *d*^−1^) the saturated soil conductivity. We neglect the effect of *ψ*_*o,soil*_ and *ψ*_*o,x*_ (the osmotic potential in the soil and in the xylem) on the flow. *ψ*_*m,soil*_ can thus be used by the xylem module (Dirichlet boundary condition), see Eqn.(29). The coupling between CPlantBox and DuMu^x^ is made available via a python binding within the root-soil interaction module DuMu^x^-ROSI [29].

We do no represent in this implementation the variation of soil carbon concentration.

#### 4 Numerical solution and computational loop

The numerical solutions of each CPlantBox modules and of the DuMu^x^ soil water flow module are described in the appendix H and in Koch et al. [31] respectively. Briefly, the soil water flow is solved by a cell-centered finite volume method (CC-FVM) [31]. The xylem water flow is solved using the hybrid analytical method of Meunier et al. [39]. The sucrose balance (sucrose usage and transport in the sieve tube and mesophyll) is computed by discretizing the continuous equation into a series of ODEs solved numerically (Newton iterations with implicit time stepping) on each plant-node (vertex-centered finite volume method, VC-FVM) [25]. The xylem water flow and FcVB-Stomatal regulation are computed together via fixed-point iterations. For these two modules, an iterative calculation of *c*_*i*_, *A*_*g*_, *g*_*co*2_, *j*_*lat,x−out*_, *ψ*_*t,ox*_, and *ψ*_*t,x*_ for each plant segment is carried out (in that order). Convergence is said to have been reached when the relative difference between two consecutive loops in each segment for each of these variables is bellow 1% or when the absolute difference is below a set threshold. The other modules are run sequentially at each user-defined time step. The user can run all the modules at each loop or increase the time step of some modules by running them every set number of loops (operator splitting). The optimal time step for the data exchange between the modules is strongly problem-dependent and to be selected by trial and error. Figure 4 gives a graphical representation of the simulation loop and main exchange of data between the modules.

**Figure 4:**
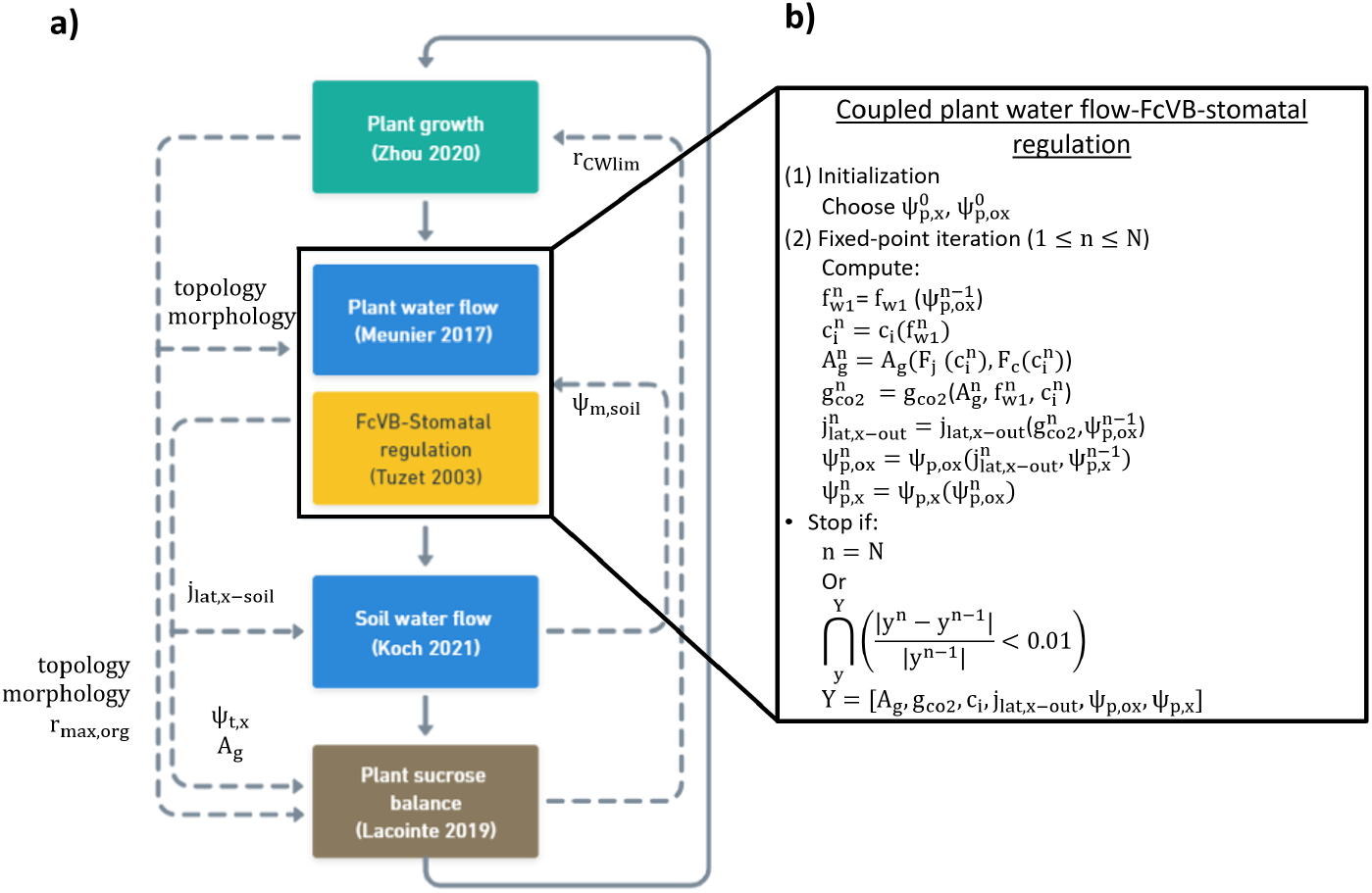
Representation of the simulation loop of the model with (a) the overall loop, and (b) the fixed-point iteration loop used for the coupled plant water flow-FcVB-stomatal regulation modules. The dashed arrows represent the exchange of data between the modules. The full arrows give the sequence of the modules in the loop. At each time step, each part of the model is computed sequentially, except for the water flow-FcVB-stomatal regulation modules which are looped over until convergence.

**Figure 5:**
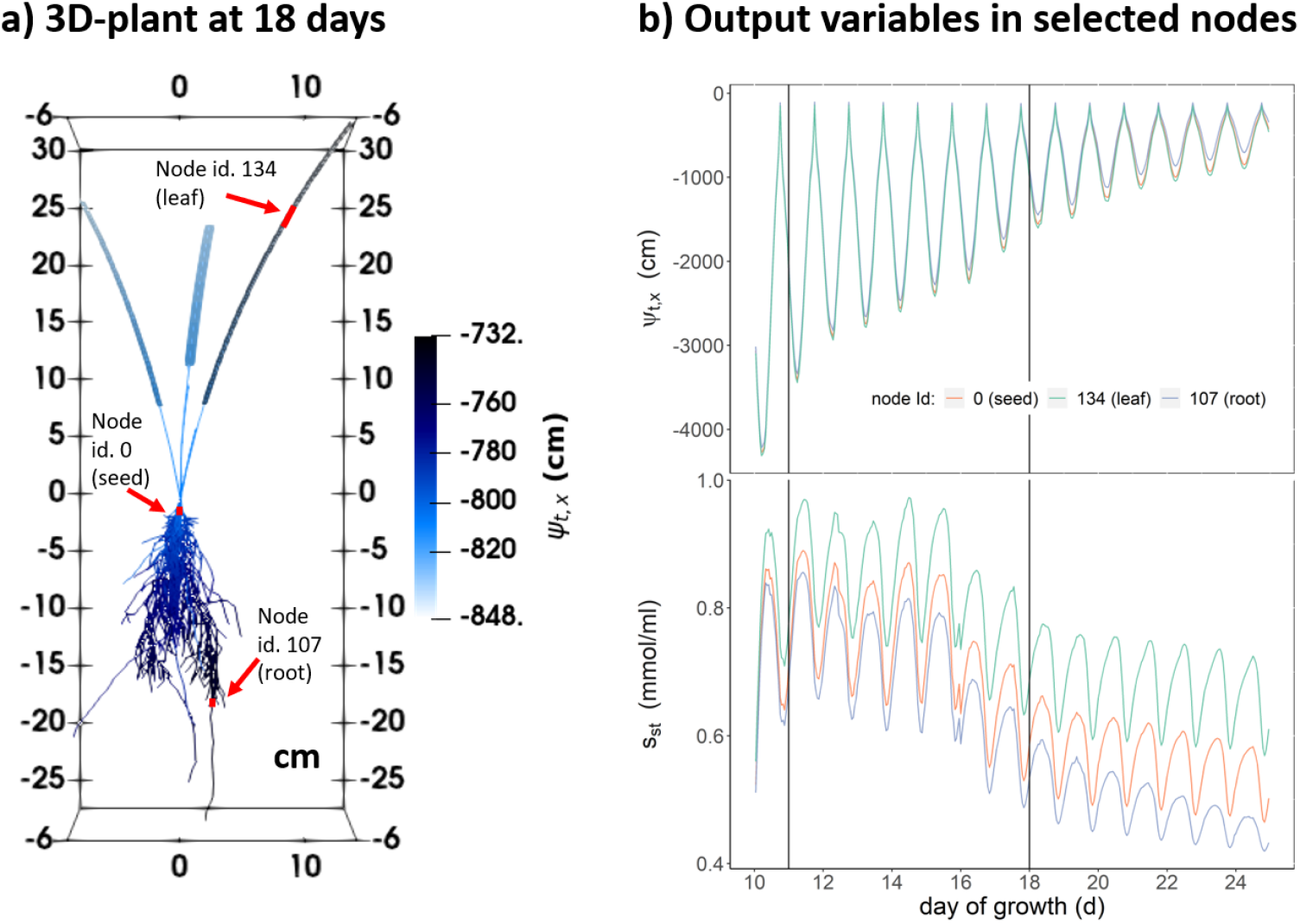
Representation of some CPlantBox outputs at the end of a simulation. a) a 3-dimensional image of a C3 monocot at 18 day under *early dry spell—wetter&colder* b) values for three specific plant nodes of the xylem water potential (*ψ*_*t,x*_) and phloem sucrose concentration (*s*_*st*_). The black vertical lines define the start and end of the dry spell (no rainfall). The red arrows and rectangles indicate the selected nodes.

**Figure 6:**
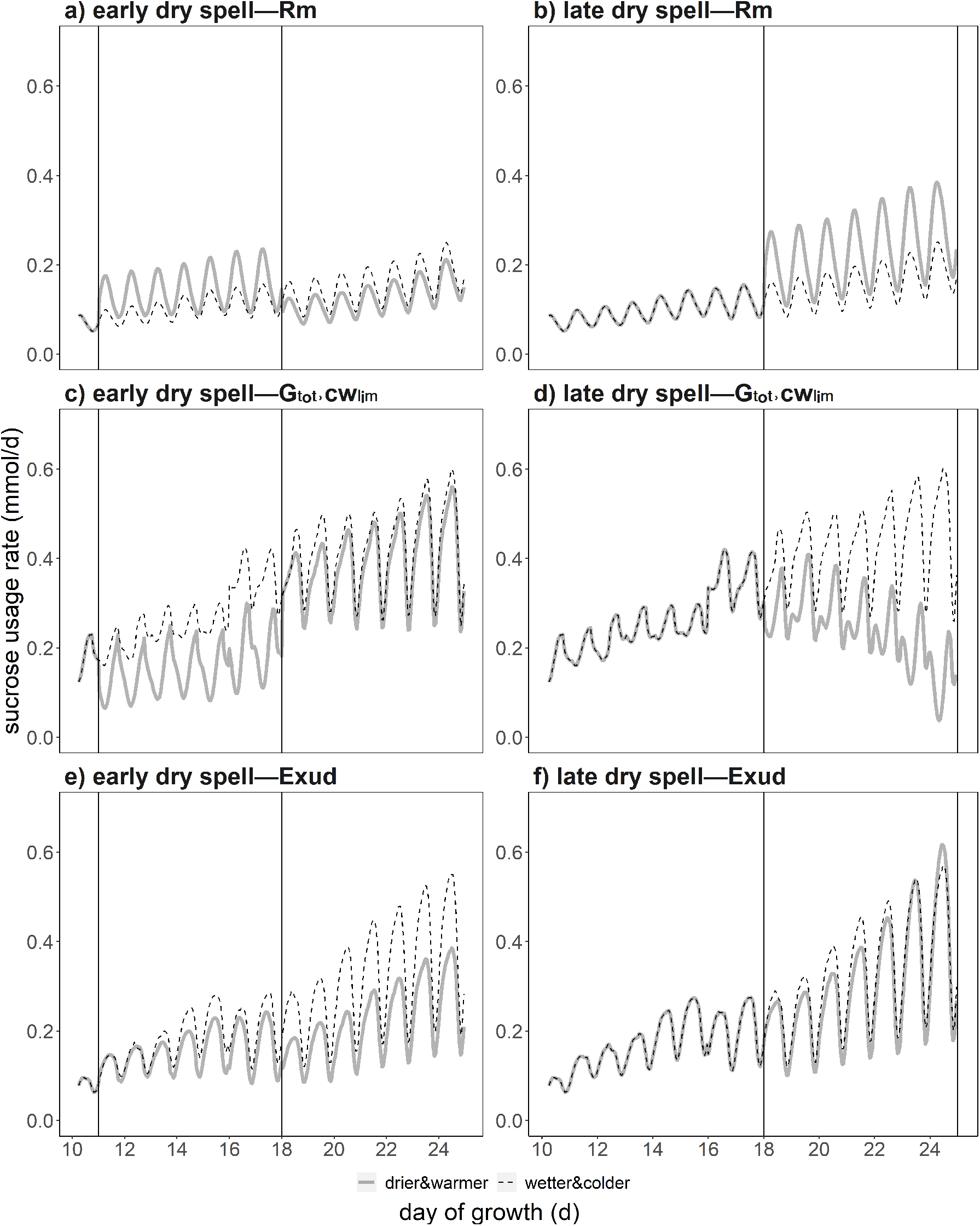
Partitioning of the carbon loss between the three sinks according to time: maintenance respiration (*Rm*), growth and growth respiration (*G*_*tot,CW lim*_), root exudation (*Exud*). The black vertical lines define the start and end of the atmospheric dry spells (no rainfall). We compare wetter and colder dry spells (thin dotted lines) against drier and warmer dry spells (thick lines).

## Part II

### Application of the model

#### 1 System modelled

The model was calibrated for C3 monocots using experimental (see Appendix G.1) and literature data. The list of the parameters and their value is given in Appendix G.2.

Notably, for this implementation, different *k*_*lat,st,root*_ and *k*_*lat,x,root*_ were defined for each root order. Moreover, we set

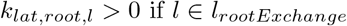

where *l*_*rootExchange*_ is between 0 and 0.8 *cm* from the root tip. This means that root water uptake and exudation are assumed to occur in the immature portion of the root [7, Chap. 4].

We simulated the effects of a dry spell of [A] two intensities [B] at two stages of development of our virtual C3 monocot.

[A] Both dry spells were represented by a period one week without rainfall. The environmental variables (see Table 1) of the low-intensity (resp. high-intensity) dry spell were set from the data of Swart et al. [59] for Germany in June of 2010 (resp. 2100, under the *SSP* 5 − 8.5 scenario), making our baseline the wetter and colder (resp. our alternative the drier and warmer) dry spell, thereafter called *wetter&colder* (resp. *drier&warmer*). *θ*_*mean,soil*_ for *wetter&colder* was set near saturation. *θ*_*mean,soil*_ for *drier&warmer* was set 30% lower, which corresponds to the relative decrease in moisture in the upper soil column between June 2010 and 2100 [59].

**Table 1:** Environmental variables.

| dry spell | min-max of sinusoidal function<sup>a</sup> | | | $\theta_{mean,soil}$ |
| --- | --- | --- | --- | --- |
| | $PAR$ | $T_{atm}$ (K) | $h_{atm}^b$ (%) | ($cm^3\ cm^{-3}$) |
| *wetter&colder* | 0-940 | 288.95-295.15 | 0.6-0.88 | 0.4 |
| *drier&warmer* |  | 293.85-303.42 | 0.44-0.74 | 0.28 |

<sup>a</sup> $T_{atm}$ , $PAR$ and $h_{atm}$ go from their minimum to their maximum value via a sinusoidal function with a period of 1 day to represent the day-night cycle. All the atmospheric variables correspond to values 2 m aboveground.
<sup>b</sup> relative air humidity.
the *wetter&colder* variables were used for all simulations before the start of the dry spells

[B] The dry spells under each scenario were simulated for two growth periods of the plants: 11 to 18 days and 18 to 25 days after sowing, hereafter called *early dry spell* and *late dry spell*.

The virtual plants at the beginning of the dry spells were obtained by running CPlantBox up to day 10 using a mean empirical growth rate (neither dependent on the carbon- nor water-flow) and without simulation of the water and carbon fluxes. Indeed, the plant leaf blades (the carbon source and water sink) had to be large enough before starting the carbon- and water-limited growth. After three hours of burn-in time, we simulated of the carbon and water fluxes and of the carbon-and water-limited growth. The plants all grew under the *wetter&colder* conditions with a static soil (constant mean *ψ*_*m,soil*_ = 0.4 *cm*^3^*cm*^−3^ at hydraulic equilibrium) between day 10 and the start of the dry spells (11d, resp. 18d). At the start of the dry spells, the dynamic soil water flow module was implemented. For the plants under *drier&warmer*, the alternative initial soil conditions was used (initial *ψ*_*m,soil*_ = 0.28 *cm*^3^*cm*^−3^). The soil water flow was simulated until the end of the dry spells (18d, resp. 25d). Therefore, at the start of dry spell period, the two *early dry spell* (resp. *late dry spell*) monocots were exactly identical. After the dry spells, the *wetter&colder* environmental and static soil variables were used. Table 2 summarises the above-given timeline of the four simulations.

**Table 2:**
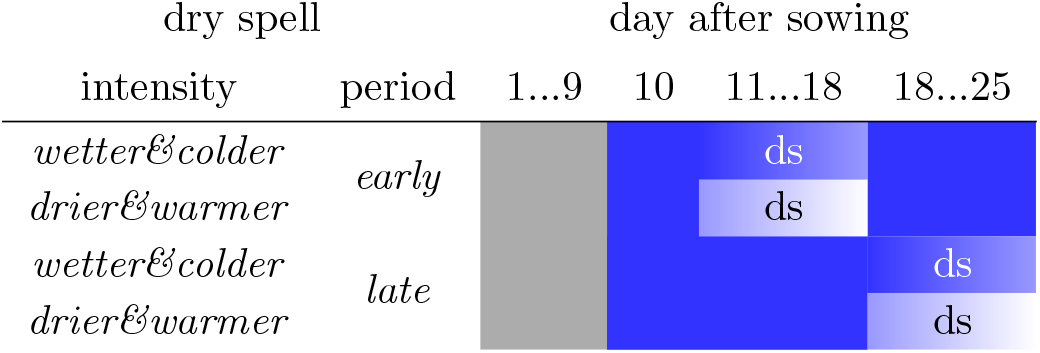
Timeline of the simulations. The grey (resp. blue) cells indicate the empirical (resp. semi-mechanistic) growth period and “ds” the implementation of the dynamic soil. The blue gradient during the dynamic soil simulation represent the decreasing soil water content. The sharp shift from dark to light (resp. light to dark) blue at the beginning (resp. end) of the dry spells for the *drier&warmer* scenario (resp. both scenarios) represent the implementation of the *drier&warmer* (resp. *wetter&colder*) atmospheric and initial (resp. constant) soil variables.

The minimum plant segment length was set to 0.5 *cm*. The *θ*_*soil*_ per voxel is (until the start of the dry spells) constant per layer and varies with the depth (hydraulic equilibrium). The mean *θ*_*soil*_ value is given in table 1. The voxel size (resolution) was set to 1 *cm*^3^.

The Van Genuchten parameters are presented in Appendix G.2 Table 6. The mean soil soluble carbon concentration was set as a unique constant value. In order to represent the effect of the water uptake by neighboring root systems, we defined the lateral dimensions of the simulation domain to be equal to the inter plant distance, 3 *×* 12 *cm*^2^, and used a laterally periodic domain. The depth of the simulation domain was 60 *cm*. During the dynamic soil simulation, no rainfall occurred and we assumed that the soil was covered with gravel (negligible evaporation). We could thus set a no-flux upper boundary condition for the soil water flow module. We moreover set a free-flow lower boundary condition.

We assumed a constant wind speed 2 *m* above ground *u*_2*m*_ of 8.467*e* +7 *cm d*^−1^ (or 2 *m s*^−1^ in SI units). The CO_2_ concentration on the leaf surface (*c*_*bl*_) was set constant at 350*e −* 6 *mmol mmol*. We simulated a unified temperature in the whole system *T*_*plant*_ = *T*_*soil*_ = *T*_*atm*_. *PAR, T*_*atm*_ and *h*_*atm*_ (relative air humidity 2 *m* aboveground) went from their minimum to their maximum value via a sinusoidal function with a period of 1 day.

The time step for the main simulation loop was of 1 *hr*. Within this loop, the time step for the exchange of data between the plant and the soil water modules was 1 minute.

## 2 Results

### 2.1 Simulation of the time-course of water and carbon flows

The model was implemented on a Lenovo Thinkpad T490, with an Intel(R) Core(TM) i5-8265U processor, using the windows subsystem linux Ubuntu20.04. The run-time of the simulation was of about 1 hour 40 minutes. The number of plant nodes went up to 2253 (for *late dry spell—wetter&colder*). The relatively high run-time of the simulation was linked to the high number of nodes and will be lower for plants with smaller root systems or when decreasing the resolution (increasing the minimum plant segment length). Moreover, the small time step for the the exchange of data between the plant and soil water flow modules (1 minute) strongly increased the simulation time. A larger this time step will lead to a shorter computation time. It is also possible to adapt the absolute and relative error tolerances of the phloem module.

Figure 5 illustrates the simulated plant structure and output by the model. Figure 5.a shows a 3-dimensional visualisation of the monocot at 18d for *early dry spell—wetter&colder*. Figure 5.b shows examples of run-time variables for selected nodes: the total water potential in the xylem (*ψ*_*t,x*_, in *cm*) and the mean concentration of sucrose in the sieve tubes (*s*_*st*_, in *mmol cm*^−3^).

We used a spin-up period of one day before starting the *early dry spell* atmospheric dry spell to give time for the plant to be at equilibrium with its environment. However, the growth of the 2^*nd*^ order laterals led to rapid changes even after this spin-up period. Indeed, we can observe a higher variation of *ψ*_*t,x*_ and *s*_*st*_ during the first days of the mechanistic growth simulation: the small root system (limited water uptake capacity) made the *ψ*_*t,x*_ very sensible to changes in the transpiration rate. Afterwards, the development of the lateral roots allow the plant to keep higher *ψ*_*t,x*_ values during the day.

### 2.2 Sucrose usage and non-structural sucrose content

Figure 6 presents the sucrose partitioning between the different sinks. Figure 14 in Appendix J presents the same data but over a shorter period to give a clearer example of the daily *s*_*st*_ variations. During the dry spells, the absolute value of maintenance respiration (*R*_*m*_) was higher for *drier&warmer* because of the higher temperature, leading to a higher enzyme activity. The daily *R*_*m*_ peak was likewise caused by the higher temperature at midday.

Because of the sink prioritisation (maintenance > growth), in case of low assimilation (night time), growth respiration (*G*_*tot,CW lim*_) diminished to fulfill the *R*_*m*_ need.

The minimum soil matric potential (*ψ*_*m,soil*_) diminished during the dry spells. At the beginning of the dry spells (18d or 25d) minimum *ψ*_*m,soil*_ (*ψ*_*m,soil*_ in the upper soil layer) was −117 *hPa* for *wetter&colder* and −433 for *drier&warmer*. At the end of the *early dry spell* dry spell simulations (at 18d) *ψ*_*m,soil*_ = −134 *hPa* for *wetter&colder* and −3824 *hPa* for *drier&warmer*. At the end of the *late dry spell* dry spell simulations (at 25d) *ψ*_*m,soil*_ = −150 *hPa* for *wetter&colder* and −5221 *hPa* for *drier&warmer. ψ*_*m,soil*_ affected the symplasm turgor potential (*ψ*_*p,symplast*_): For *early dry spell*, the minimum *ψ*_*p,symplast*_ during the dry spell was of 3532 *hPa* and 6575 *hPa* for *drier&warmer* and *wetter&colder* respectively. For *late dry spell*, the minimum *ψ*_*p,symplast*_ during the dry spell was of 2663 *hPa* and 8427 *hPa* for *drier&warmer* and *wetter&colder* respectively. For *drier&warmer, ψ*_*p,symplast*_ was nearer the critical symplasm turgor potential for growth below which no growth occurs (*ψ*_*p,crit*,2_, set to 2000 *hPa*), leading to a lower *G*_*tot,CW lim*_. Water availability had therefore a strong effect on *G*_*tot,CW lim*_.

*Exud* varied according to *s*_*st*_ and was lower at night time. Moreover, we saw diverging variations of *Exud* during the dry spells. Indeed, the lower *G*_*tot,CW lim*_ for *early dry spell—drier&warmer* led to a slower growth of the leaf blades (sucrose source) and of the root system (exudation zone), causing the lower daily maximum *Exud* for *drier&warmer* when compared with *wetter&colder*. The difference became higher after the end of the dry spell. For *late dry spell—drier&warmer*, the higher *R*_*m*_ at the beginning of the spell (due to the higher temperature) caused a lower *s*_*st*_ (see Figure 7). Near the end of the spell, the stronger decrease in *G*_*tot,CW lim*_ compensated the *R*_*m*_ increase and led to a higher maximum daily *Exud*.

**Figure 7:**
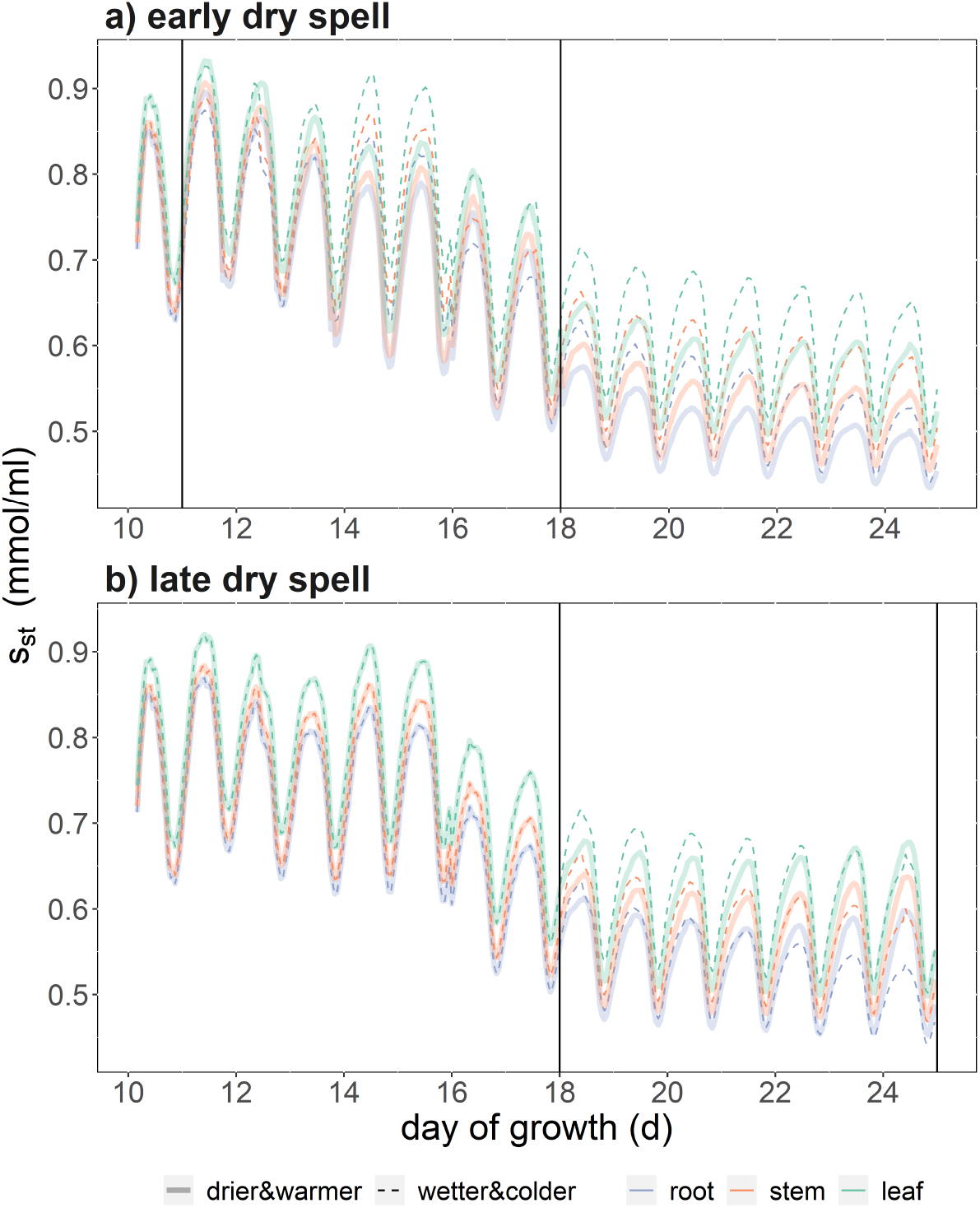
Mean sucrose concentration (*s*_*st*_) according to time in the root (blue), stem (orange), and leaf (green) organs. The black vertical lines define the start and end of the dry spells (no rainfall). We compare wetter and colder dry spells (thin dotted lines) against drier and warmer dry spells (thick lines).

At the end of the simulation, *early dry spell—wetter&colder* and *late dry spell—wetter&colder* have similar cumulative *G*_*tot,CW lim*_, *R*_*m*_, and *Exud*. Compared with *wetter&colder, early dry spell —drier&warmer* led to a strong decrease of cumulative *G*_*tot,CW lim*_ (−27%), and *Exud* (−26%) and to a slight increase of cumulative *R*_*m*_ (+4%). *late dry spell—drier&warmer* led to a strong decrease of cumulative *G*_*tot,CW lim*_ (−36%), a slight decrease of cumulative *Exud* (−6%) and to a strong increase of cumulative *R*_*m*_ (+37%).

Figure 7 presents the mean *s*_*st*_ per organ type. During the dry spell of the *early dry spell* simulation, there was a compensatory effect of the higher *R*_*m*_ and lower *G*_*tot,CW lim*_ and *s*_*st*_ was not strongly different between *drier&warmer* and *wetter&colder*. After the dry spell, the lower leaf growth led to a lower total assimilation and *s*_*st*_ became lower for *drier&warmer* compared with *wetter&colder*. For *late dry spell*, the stronger decrease in *G*_*tot,CW lim*_ led to a higher *s*_*st*_ for *drier&warmer* compared with *wetter&colder*. This differences in *s*_*st*_ in the root caused the different *Exud* observed in Figure 6.

### 2.3 Structural sucrose content

Figure 8 presents the absolute structural sucrose content for each organ type and subtype.

**Figure 8:**
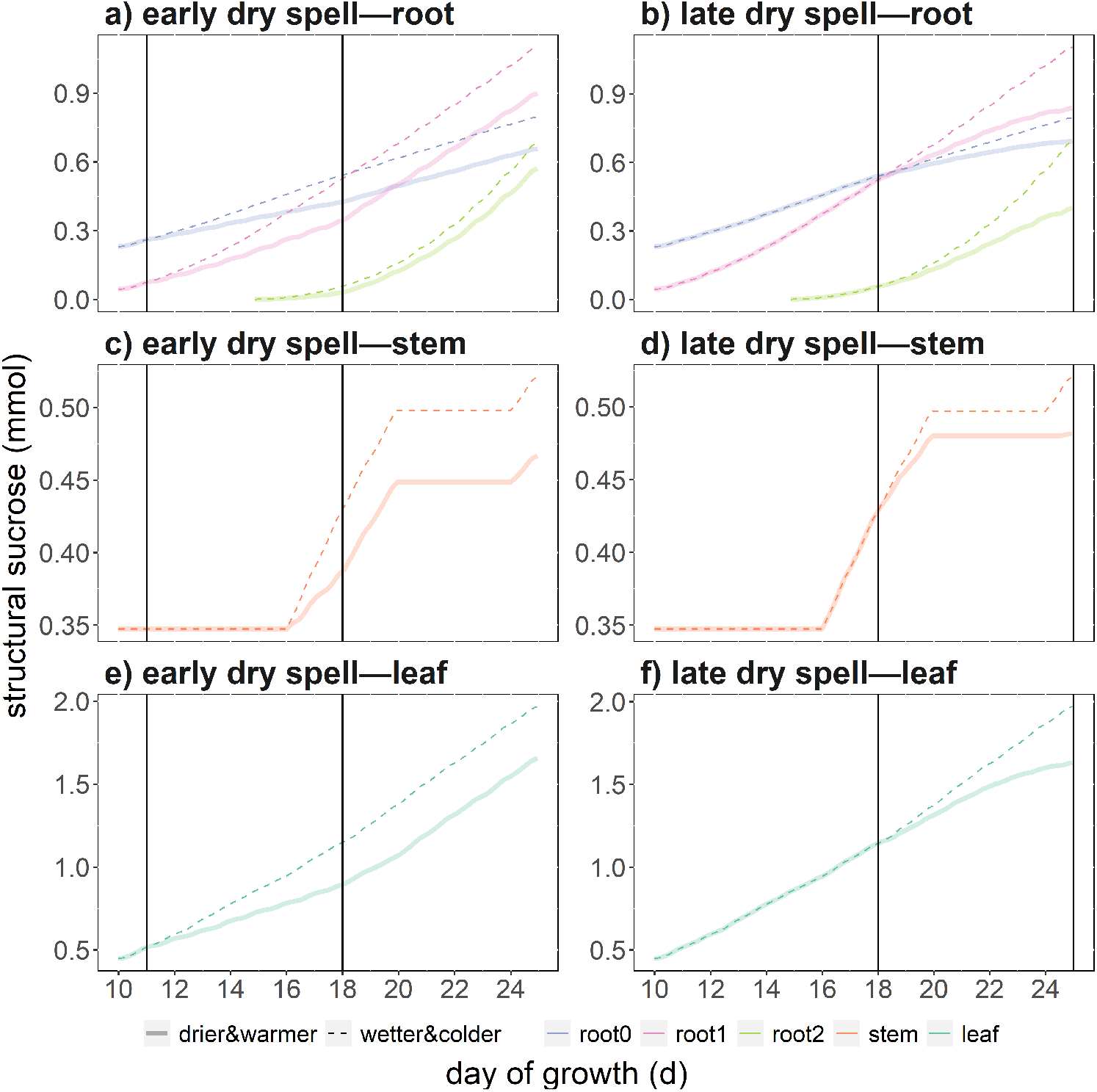
Absolute structural sucrose content per plant organ type and subtype according to time, with 0-order roots (root0, blue); 1^*rst*^ order roots (root1, pink), 2^*nd*^ order roots (root2, light green), stem (orange), and leaf (dark green) organs under drier and warmer conditions (thick transparent line) or wetter and colder conditions (thin lines) for two development stages of the plant. The black vertical lines define the start and end of the dry spells (no rainfall). We compare wetter and colder dry spells (thin dotted lines) against drier and warmer dry spells (thick lines).

At the beginning of each dry spell (11d or 18d) the monocots under *drier&warmer* and *wetter&colder* are exactly identical. We can observe several growth plateaus in the developme It was used to represent the period after the first appearance of the stem and before the stem elongation phase.

Moreover, the lower *G*_*tot,CW lim*_ under *drier&warmer* (see Figure 6) led to a lower growth for all organ types. Especially, we see a lower growth rate for *late dry spell—drier&warmer* compared with *drier&warmer* during the last three days of the simulation, which fits with the lower *G*_*tot,CW lim*_ observed in Figure 6. For all simulation scenarios, we can observe that the growth rate of the 2^*nd*^ order laterals is much higher after day 18, during the later dry spell.

Figure 9 presents the relative structural sucrose content for each organ type and subtype.

**Figure 9:**
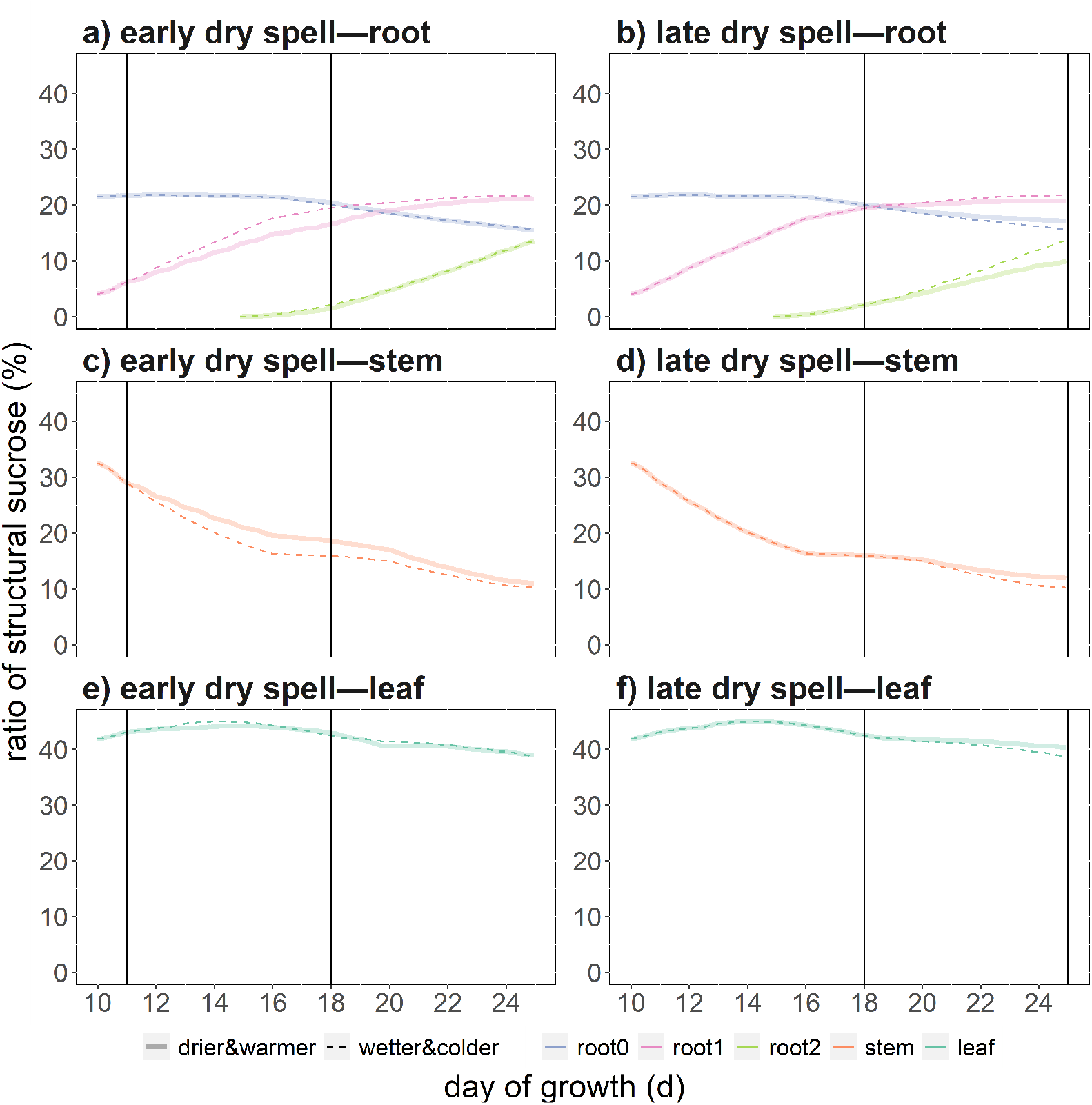
Relative structural sucrose content per plant organ type and subtype according to time, with 0-order roots (root0, blue); 1^*rst*^ order roots (root1, pink), 2^*nd*^ order roots (root2, light green), stem (orange), and leaf (dark green) organs under drier and warmer conditions (thick transparent line) or wetter and colder conditions (thin lines) for two development stages of the plant. At the start of each dry spell (11d or 18d) the monocots are identical between the two weather scenarios.

For all simulations, the root-to-shoot ratio of structural sucrose content increased during the simulation. We observed that the dry spell intensities had a slight effect on the relative structural sucrose partitioning.

For instance, the relative sucrose content in the root was of 50% for *early dry spell—drier&warmer*, 51% for *early dry spell—wetter&colder*, 48% for *late dry spell—drier&warmer*, 51% for *late dry spell—drier&warmer*.

Because the growth of the 2^*nd*^ order roots was much more important after day 18, *late dry spell—drier&warmer* had the lowest relative structural sucrose content of 2^*nd*^ order roots (10%) when compared with *early dry spell—drier&warmer* (13%) or *wetter&colder* (14%).

Figure 10 shows the 3D representation of the plants at the end of the simulation. Under *wetter&colder* (10.a and 10.c) we observed a higher development of the leaves and root systems compared to *drier&warmer* (10.b and 10.d). Indeed, the lower increase in root structural carbon is reflected by the different total root lengths per root order at day 25: for *early dry spell*, we obtained 2460 *cm* against 2985 *cm* for *drier&warmer* and *wetter&colder* respectively. For *late dry spell*, we obtained 2104 *cm* against 3003 *cm* for *drier&warmer* and *wetter&colder* respectively.

**Figure 10:**
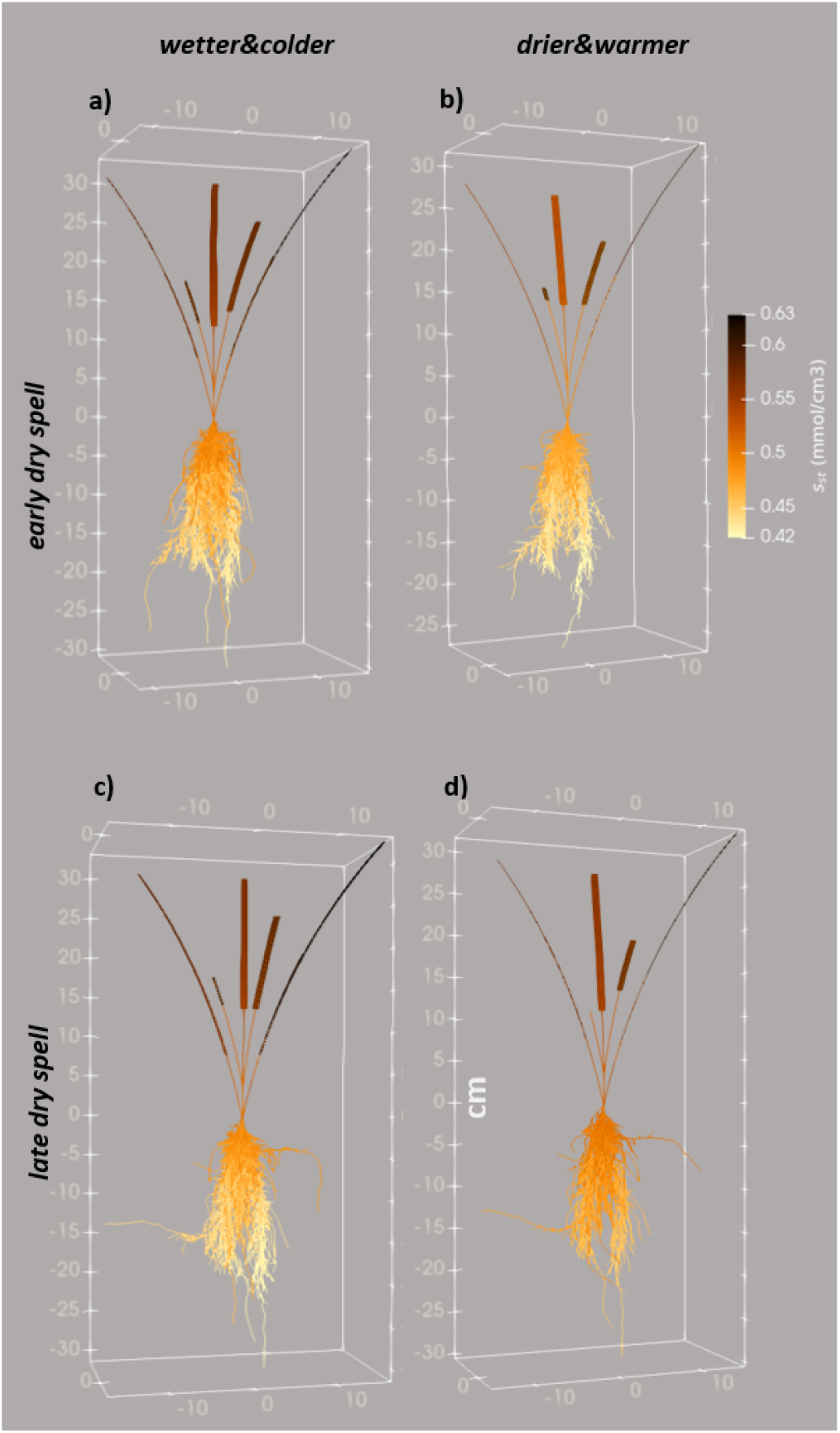
3D representation of the virtual plants at 25d after a a) wetter and colder dry spell between days 11 and 18, b) drier and warmer dry spell between days 11 and 18, c) wetter and colder dry spell between days 18 and 25, d) drier and warmer dry spell between days 18 and 25. Each segment is colored according to its sucrose concentration in the sieve tube (*s*_*st*_, *mmol cm*^−3^).

Likewise, we can observe the different total leaf length per plant at day 25: for *early dry spell*, we obtained 139 *cm* against 159 *cm* for *drier&warmer* and *wetter&colder* respectively. For *late dry spell*, we obtained 137 *cm* against 160 *cm* for *drier&warmer* and *wetter&colder* respectively.

Moreover, Figure 10 allows us to see the *s*_*st*_ gradient along the plant’s organs. For instance, longer mature leaves had the highest *s*_*st*_ values.

### 2.4 Instantaneous water use efficiency

Figure 11 presents the cumulative transpiration and sucrose assimilation (*A*_*g*_) for both dry spell scenarios (*drier&warmer, wetter&colder*) and periods (*early dry spell* and *late dry spell*).

**Figure 11:**
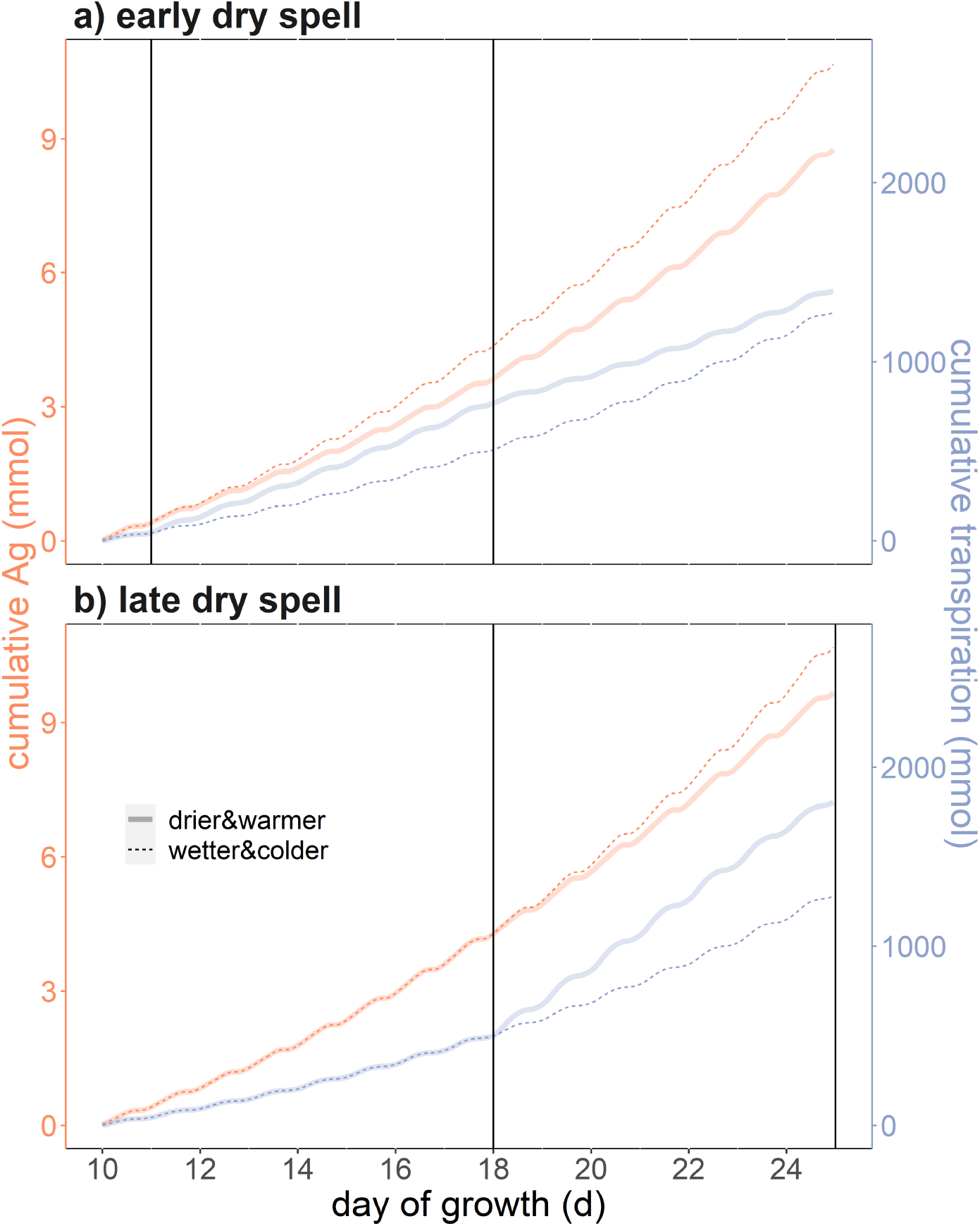
Cumulative assimilation of sucrose (*A*_*g*_, in orange) and cumulative transpiration (blue) according to time under drier and warmer conditions (thick transparent line) or wetter and colder conditions (thin lines) for two development stages of the plant.

*drier&warmer* had a negative effect on cumulative *A*_*g*_: −18 % for *early dry spell* and −9 % for *late dry spell*. this is in part caused by the lower leaf development (see Section 2.3).

For all four simulations we observed a closing of the stomata because of a low leaf water potential near midday. During the dry spells, the water scarcity factor for stomatal opening (*f*_*w*1_) went down to 0.82 for both *wetter&colder* scenarios, 0.64 for *early dry spell—drier&warmer* and 0.55 for *late dry spell—drier&warmer*. A smaller opening of the stomata led to a lower substomatal CO_2_ concentration (*c*_*i*_) and through this also caused the lower *A*_*g*_ for *drier&warmer*. Moreover, the higher leaf to air vapor pressure deficit during the dry spell led to a higher cumulative transpiration for *drier&warmer* compared with *wetter&colder* : +20% for *early dry spell,drier&warmer* and +40% for *late dry spell,drier&warmer*.

Overall, we observed a decrease in instantaneous water use efficiency (*Ag*-to-transpiration ratio) at the end of the simulation for *drier&warmer* compared with *wetter&colder* : −31% for *early dry spell* and −35% for *late dry spell*.

## Part III

### Discussion

#### 1. CPlantBox offers a user-friendly flexible representation of the interactions between plant development and water- and carbon-fluxes

In this model, we implemented modules which were already included in other FSPMs: 3D representation of shoots and roots [55, 67], linked carbon and water fluxes [55, 67, 32], carbon respiration [3, 22], exudation [67], distributed carbon sources and sinks [32], water and/or carbon dependent growth [46, 13, 36, 22], coupled soil and root water flux [29], effects of the atmosphere variables and plant water status on the coupled stomatal opening and photosynthesis [64]. The novel aspect of this latest implementation of CPlantBox comes from linking all of these modules within one single user-friendly framework [36] to represent the carbon and water fluxes in the soil-plant-atmosphere-continuum. This allowed us to look at the feedback effects between plant growth and water- and carbon-fluxes, which was lacking in earlier models [14, 57, 16, 67]. Processes which are usually predefined in FSPMs, like carbon partitioning [46, 57] became consequently emerging properties of our model. We could therefore simulate how growth and carbon partitioning vary according to the growth conditions via semi-mechanistic process descriptions. CPlantBox is also a flexible model: the spacial and temporal resolution (respectively plant segment length and time step) can be defined at run-time by the user. CPlantBox can also represent a wide variety of plants, contrary to other available FPSMs [13, 36].

This is in agreement with the aims of CPlantBox which is to be a tool to [a] test the effect of the interactions between plant genotype, environmental conditions, and agricultural management, [b] help understand how/why these interactions lead to emerging properties (like plant structure and photosynthetic capacity), [c] show as output parameters used as inputs by crop models (like carbon partitioning).

On the computing side, CPlantBox represents organisms using graph formalism (similarly to OpenSimRoot [46]). This representation allows us to implement existing solvers created for graphs or 1-dimensional meshes [25, 23, 31]. These solvers greatly increase the efficiency of the flow computation [32]. Likewise, the implementation of CPlantBox in C++ with Python binding allows us to use the rapidity of C++ and the flexibility and clarity of Python. This is a non-negligible advantage as computational power becomes a significant limitation for the development of FSPMs [57, 13].

### 2 Results of the simulation

In this study, we used our model to look at the effect of a dry and warm spell of one week on plant water flow, carbon flow, and growth, based on the expected temperature, air relative humidity, and relative variation in soil water content in Germany for June of 2100 (but without changes of CO_2_ concentrations).

We could observe a lower growth rate for all organ, fitting with the results of Chen et al. [10] for monocots under drought stress. In this simulation, the lower growth was caused by a too low turgor pressure rather than a lack of sucrose.

We observed a lower assimilation-to-transpiration ratio under the drier and warmer scenario, which was also observed experimentally by Tshikunde et al. [63] for a monocot. This strong influence of the leaf area on the plant’s photosynthetic capacities confirms the assumption of Damour et al. [14] in their review of stomatal regulation models.

During the simulation, the ratio of exudation to total carbon usage remained within the range found in the literature for monocots: 3 − 40% [19]. However, as soil soluble carbon concentration was a constant during the simulation, we could not model the feedback effects of different exudation levels. The plant sucrose concentration varied between 0.4 *mmol ml*^−1^ and 0.93 *mmol ml*^−1^, which fits with the values reported by [60, 27]. The low pressure gradient between source and sinks follows as well the current assumption for herbaceous angiosperms [15].

Contrary to other modelling studies, the drier and warmer scenario did not lead to a decrease in transpiration [64, 4]. Indeed, the slight closing of the stomata did not compensate the higher evaporative demand caused by the lower air relative humidity. Moreover, the initial soil water potential remained high. The chosen grid resolution for the soil (1 *cm*^3^) was also relatively coarse and might have led to an over-estimation of the plant water uptake and thus of the transpiration [29]. Moreover, we did not represent the loss of conductivity caused by cavitation and this process can strongly affect the transpiration rate [12, 16]. Likewise we did not simulate the stomatal closure linked to an increase in *s*_*meso*_ [15], which would have lead to a lower assimilation and transpiration for the plants at the beginning of the simulation, when *s*_*st*_ is higher. Divergences between our model and other observed or simulated results remain however useful as they can give insight into which processes are key to representing emerging properties.

### 3 Current limitations of the model and future perspectives

The increasing number of modules in CPlantBox leads to a high number of parameters and (intermediary) outputs. Also, several variables and parameters cannot be measured experimentally (like the separation of the respiration between maintenance and growth [61]) making their experimental evaluation more complex.

A first sensitivity analysis (see appendix I) was therefore done to evaluate the most important parameters. Moreover, several controls were setup in CPlantBox to warn the user in case of impossible outputs (like negative transpiration rate and too high sucrose concentration in the sieve tube). Building on this work, the development of an effective calibration pipeline and parameter database would be an important step towards making this model more readily usable [36]. On the experimental side, new (field) experiments will be conducted to parameterize the model for a specific wheat genotype rather than a generic C3 monocot. When CPlantBox is not precisely parameterized, it can be used to do qualitative tests rather than quantitative predictions.

The fixed-point approach used for the plant water flow-photosynthesis-stomatal regulation modules (Sections 2.2 and 2.1) always converged for our simulations (convergence reached in *<* 5000 loops). Including other modules in the fixed-point iteration (like the soil water flow module) would make the model outputs less dependent on the selected time step.

Regarding the processes represented, the accuracy of the phloem flow outputs could be further improved by adapting the representation of the sucrose usage. First of all, the plant’s respiration is currently following the widely-used growth-maintenance paradigma. As other processes/functional modules are added into CPlantBox (like nitrogen uptake and flow), it will be possible to shift to a process-based definition of the respiration rate, as recommended by Cannell and Thornley [9] and implemented by Gauthier et al. [22]. This would also improve the evaluation of the feedbacks between the environmental conditions and the plant’s processes. Also, the representation of the sucrose exudation rate, and of the chlorophyll-dependent assimilation offers the first elements of a more explicit representation of the nutrient flow within the plant, which are an important aspects of a plant’s adaptation to its environment [21]. Moreover, the representation of [a] reversible soluble sugar to starch conversion and [b] sucrose-induced closing of the stomatas would help evaluate the plant’s sucrose-related adaptation capacity to a changing environment [15]. Indeed, starch synthesis and hydrolysis can help the plant regulate sucrose concentration and withstand periods of low assimilation. The negative feedback of the sucrose on photosynthesis would on the other hand help better represent the decrease of photosynthetic capacity in case of low sucrose usage. Representing tissue death caused by water and/or carbon scarcity would allow CPlantBox to simulate plant development under more extreme weather conditions. Finally, the explicit 3-dimensional representation of the shoot organs makes it possible to simulate precisely the atmospheric variables (like *PAR, T, g*_*bl*_, *g*_*canopy*_) for each leaf segment. Next implementations of the model will simulate micro-climate in the canopy.

## Conclusion

In this paper, we presented the latest version of CPlantBox and implemented it to represent the effect of dry spells of one week, starting at different days (11 against 18 days after sowing) and with different intensities (wetter and colder against drier and warmer). We could observe that the drier environment led to higher respiration and exudation rates. The model also showed that, under the warmer and drier climate, the early dry spell affected the plant processes (higher transpiration, lower assimilation, lower organ growth) less than the late dry spell when compared with the wetter and colder climate.

The model was therefore capable of computing plant variables at a sub-diurnal scale and for each plant control volume in a complex architecture (fully grown monocot with distributed water and carbon sink and sources). The model could moreover simulate semi-mechanistically the interplay between the three dimensional plant topology and the environmental conditions via the water and carbon flows in the soil-plant-atmosphere continuum by representing a growing plant in a dynamic environment. The computed variables at small time and spatial scale led to emerging property which are usually empirically defined in plant models (growth rate, carbon partitioning).

CPlantBox is a versatile open-source tool which can be used to test hypotheses, select experimental setups, guide genotype selection, and predict and understand qualitative variations of observed experimental results under different genotype-environment-management combinations.

Next steps in the development of CPlantBox will include the setup of a calibration pipeline and the gathering of field and laboratory data to represent a specific plant phenotype.

## Data and code availability

The results of the experiments are available upon request. The code used to run the simulations is available at https://github.com/Plant-Root-Soil-Interactions-Modelling/CPlantBox/releases/tag/v2.0.

## Acknowledgement

The authors thank S. Moyo, M. Humza, C. Tempelmann, E. Vogt and A. Hollweg for their technical assistance during this work

## Author contributions

SLG, YR, and MG conceived the experiments. SLG and MH ran the experiments under the supervision of YR and DvD. MG implemented the model changes described in Appendix A under the supervision of AS, GL, DL and JV. SLG and MG calibrated the model. MG ran the simulations under the supervision of AS and GL. DL, AS, FM, and MJ set up the equations (and wrote) Part I.2.2 and Appendix H.2.1. SL wrote Appendix G.1. MG wrote the other parts of the manuscript under the supervision of AS and GL. All authors provided critical feedback and helped shape the research, analysis and manuscript.

