## Appendices for "Development and calibration of the FSPM CPlantBox to represent the interactions between water and carbon fluxes in the soil-plant-atmosphere continuum"

#### A comparison with the former CPlantBox implementation

The functional-structural plant model CPlantBox was first presented by Zhou et al. [67]. The differences between the version of Zhou et al. [67] and the current version of the model are presented in Figure 12.

In summary, the following modules were added or adapted compared with the original model: (a) water- and carbon-limited growth was implemented (see Section 2.2); (b) the pre-existing xylem water flux analytical solver ([39]) was expanded from the root to the whole plant and implemented (see Section 2.2); (c) the implicit numerical solver PiafMunch [32] for phloem flows driven by hydrostatic pressure gradients was adapted for tight coupling to CPlantBox (see Section 2.3); (d) the DuMu$^x$ PDE solver [31] simulating soil water fluxes with a cell-centered finite volume method (see Section 3); and (e) a coupled photosynthesis (FcVB)-stomatal opening [64] were added (see Section 2.1).

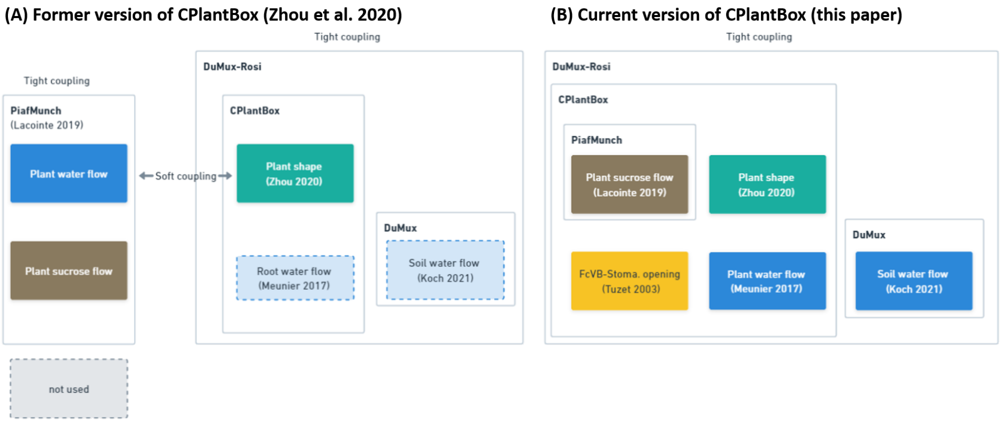


Figure 12: **Evolution of CPlantBox within the DuMu$^x$-ROSI interaction framework.** Representation of (A) the former and (B) current version of CPlantBox. Main changes include: an expansion of the root water flow module to shoot organs, implementation of a photosynthesis (FcVB) and stomatal opening module, adaptation and implementation within CPlantBox of a phloem flow module. The tighter coupling between the phloem and plant module allows us to use the CPlantBox water module and the DuMu$^x$ soil flow module.

#### B Water potentials, conductivities, and conductances

We evaluate the water potentials in four domains: the soil, the xylem, the sieve tubes, and the atmosphere, respectively denoted with the subscripts $_{soil}$, $_x$, $_{st}$, and $_{atm}$. For clarity, we define here the total potential $\psi_t$ and its components in each of those domains. We define $\psi_t$ as

$$\psi_t \quad = \quad \frac{\eta - \eta_{ref}}{\bar{V}} \quad = \quad \psi_p + \psi_g + \psi_o,$$

where $\eta$ and $\eta_{ref}$ are the chemical potentials of the considered and reference water respectively, $\psi_p$ is the pressure potential, $\psi_g$ is the gravitational potential and $\psi_o$ is the osmotic potential [43]. $\psi_p$ includes all the potentials except the osmotic and gravitational ones and can be positive (absolute $\psi_p$ > absolute $\psi_{p,ref}$) or negative (absolute $\psi_p$ < absolute $\psi_{p,ref}$). In unsaturated soils,

$$\psi_{p,soil} = \psi_{m,soil} + \psi_{a,soil} < 0$$

with $\psi_m$ the matric potential and $\psi_a$ the pneumatic potential. The pneumatic potential corresponds to the over-pressure of the air: $\psi_a = P_{atm} - P_{soil\ air}$. $P_{atm}$ ($hPa$) is the atmospheric pressure at ground level, dependent on the pressure of all the gases in the Earth's atmosphere. The reference water (denoted with the subscripts $_{ref}$) is pure, at ground level ($z_{ref} = 0$), and the hydrostatic pressure equals the atmospheric pressure. In the plant, $\psi_p = \psi_h$, with $\psi_h$ the hydrostatic potential. $\psi_p$ is generally positive in the plant tissues and is also called the turgor pressure. For specific cells, like the xylem $\psi_{p,x}$ is usually under tension. All the water potentials are given in $hPa$ (corresponding to $100 \times Joule/m^3$).

In this paper, the flow of fluids considered can be computed with either the lateral conductivity ($k_{lat,Y}$, in $L\ P^{-1}T^{-1}$, here in $cm\ hPa^{-1}\ d^{-1}$), or the intrinsic axial conductance ($K_{ax,Y}$, in $L^4P^{-1}T^{-1}$, here in $cm^4\ hPa^{-1}\ d^{-1}$) [39]. $_{lat}$ stands for lateral and $_{ax}$ for axial. $_Y$ is a stand in for the denomination of the flow channel (compartment, tissue, membrane...) considered.
For the photosynthesis module, we use the stomatal conductance in molar units ($g_{Z,Y}$, in $N\ L^{-2}T^{-1}$, here in $mmol\ cm^{-2}\ d^{-1}$), with $_Z$ standing in for either $_{h2o}$ or $_{co2}$.
For the soil module, the hydraulic conductivity ($\kappa_{soil}$), units are in $cm^2\ hPa^{-1}\ d^{-1}$ ($L^2\ P^{-1}T^{-1}$) given the definition of the soil water potential.

### C Computation of the plant shape variable

An organ's length is equal to the sum of the length of its segments: $L_{seg} \leq L_{org}$, with $L$ in $cm$. Root and stem organs are defined as cylinders, which shapes are computed from their lengths and radii ($a$ in $cm$). In the current CPlantBox implementation, $a$ is constant for all segments of the organ: $a_{seg} = a_{org}$.
Leaf organs are defined as cuboids which shapes are computed from their thicknesses ($a$ in $cm$) and widths ($Wi$ in $cm$). The leaf organs are separated in two regions: the sheath and the blade. $a$ and $Wi$ of the sheath can be different to that of the blade.
The seed is represented by one node linking the main stem (and tillers) with the 0-order roots. The sieve tubes and xylem tissues are defined as cylinders. Each organ can have one or more sieve tube or xylem cylinders. Finally, the shape of the mesophyll is assumed to be equal to that of the leaf blade it is in (for instance with $Vol$, the volume in $cm^3$: $Vol_{meso} = Vol_{blade}$).

$A_{cross}$ (cross-sectional area, $cm^2$), $Vol$ (volume, $cm^3$), and $Per$ (perimeter of the exchange surface, $cm$) of all the organs, segments, sieve tubes, and xylem tissues are calculated thus:

$$Vol_Y = \begin{cases} Vol_{sheath} + Vol_{blade} & \text{for} \quad \text{leaf organs} \\ A_{cross,Y} \times L_Y & \text{else} \end{cases} \tag{59}$$

$$A_{cross,Y} = \sum_{i}^{n_X} A_{cross,Y,i} \tag{60}$$

$$Per_Y = \sum_{i}^{n_X} Per_{Y,i} \tag{61}$$

$$A_{cross,Y,i} = \begin{cases} Wi_Y a_Y & \text{for} \quad \text{leaf blades and sheaths} \\ \pi a_{Y,i}^2 & \text{else} \end{cases} \tag{62}$$

$$Per_{Y,i} = \begin{cases} 2W_i & \text{for leaf blades and sheaths} \\ 2\pi a_{Y,i} & \text{else} \end{cases} \tag{63}$$

with $_Y$ as a stand-in for either $_{st}$ (sieve tube tissues), $_x$ (xylem tissues), $_{seg}$ (segment), $_{org}$ (organ), $_{sheath}$ (leaf sheath), and $_{blade}$ (leaf blade). $_i$ represent one cylinder of the tissue type in the studied segment. $n_Y$ corresponds to the number of tissue unit of segment (or vessel) per cross section of segment. Thus $n_{seg} = 1$. In our parameterised model, $n_{st}$ (resp. $n_x$) corresponds to the number of vascular bundles per cross-section times the number of sieve (resp. xylem) tubes of a specific radius per bundle. As $Per$ corresponds to the perimeter of the lateral exchange surface, we only take into account the width of the leaf and not its thickness, as, for our virtual plant, the stomatas are located on the upper and lower side of the leaf blade.

### D Air conductance

We separate the area outside the leaf in three zones: the laminar layer directly outside the leaf (boundary layer), the turbulent layer between the leaf boundary layer and the canopy induced zero displacement height, the turbulent layer between the canopy induced zero displacement height and the point of measurement. The effect of each layer is represented by a resistivity to the water vapor flow, respectively: $Res_{h2o,bl}$, $Res_{h2o,canopy}$, and $Res_{h2o,atm}$, all in $d\ cm^{-1}$.

$Res_{h2o,atm}$ is computed according to Allen et al. [1]:

$$Res_{atm,h2o} = \frac{ln\left[\frac{z_m - \frac{2}{3}h_p}{0.123 \times h_p}\right] \times ln\left[\frac{z_m - \frac{2}{3}h_p}{0.1 \times h_p}\right]}{k_{Karman}^2 u_z} \tag{64}$$

with $z_m$ ($cm$) height of wind and humidity measurements, $h_p$ ($cm$) the height of the highest plant node, $k_{Karman}$ (0.41 $-$) the von Karman's constant, $u_z$ ($cm\ d^{-1}$) wind speed at height $z_m$. $\frac{2}{3}h_p$ is the canopy induced zero displacement height.

We set the mean canopy resistance as [45]:

$$Res_{h2o,canopy} = \frac{\frac{1}{3}h_p}{k_{eddy,canopy}} \tag{65}$$

with $\frac{1}{3}h_p$ the mean length of the canopy turbulent layer between the leaf boundary layer and the canopy induced zero displacement height, $k_{eddy,canopy}$ ($8.64e8\ cm^2 d^{-1}$) the mean eddy covariance value, corresponding to the value given by Nobel [45] for the space 1 $m$ below the top of a maize canopy. And, assuming a constant leaf boundary layer thickness of 0.03 $cm$, we obtain

$$Res_{h2o,bl} = \frac{0.03}{Diff_{h2o}} = 1.5e-6$$

with $Diff_{h2o} \approx 2e3\ cm^2 d^{-1}$, the diffusion coefficient of water vapor in air [44].
We can then convert the physical conductances ($\frac{1}{Res_{h2o,Y}}$ in $d\ cm^1\ cm^{-2}$) in molar conductances ($g_{h2o,Y}$ in $mmol\ cm^{-2}\ d^{-1}$) [44, 64]:

$$g_{h2o,Y} = \frac{P_{atm}}{Res_{h2o,Y} \times RT_Y} \quad (66)$$

### E Maximum and carbon and water limited growth rate

In CPlantBox, we do not account for reversible (elastic) tissue deformation. $r_{max,org}$ ($cm^3\ d^{-1}$) is the maximum growth rate of the organ's growing zone when the cell's symplast are at full turgor and the xylem is at the reference water potential ($\psi_{t,x} = 0$). To compute from this the carbon need for growth ($Gtot_{max}$, $mmol\ sucrose\ d^{-1}$), we first divide this growth demand between the nodes of the segments within the growing zone ($f_{length}$). The water limited growth rate ($r_{Wlim,i}$, in $cm^3 d^{-1}$) is computed thus for the node $i$:

$$r_{Wlim,i} = r_{max,org} \times f_{w2} \times f_{length} \quad (67)$$

$$f_{w2} = \frac{max(\psi_{p,symplast}, \psi_{p,crit,2}) - \psi_{p,crit,2}}{-\psi_{o,symplast} - \psi_{p,crit,2}} \quad (68)$$

$$r_{max,org} = \frac{Vol_{t_{end},org} - Vol_{t_{init},org}}{t_{end} - t_{init}} + \frac{Vol_{t_{end},org,child} - Vol_{t_{init},org,child}}{t_{end} - t_{init}} \quad (69)$$

$\psi_{p,symplast}$ ($cm$) is the turgor water pressure within the symplast of the cell. $\psi_{p,crit,2}$ is the minimum symplast turgor potential below which no growth occurs (the "wall yield threshold"). We assume that the aploplast has the same water potential values as the xylem tissues and that the permeability of the membrane between the apoplast and the symplast is high: $\psi_{t,symplast} = \psi_{t,apoplast} = \psi_{t,x}$ [16, 54]. We thus obtain $\psi_{p,symplast} = \psi_{t,x} - \psi_{o,symplast}$. Also there is a strong osmotic regulation in the symplast [54]. Therefore, we consider that the osmotic water potential in the symplast $\psi_{o,symplast}$ is constant. $f_{w2}$ increases consequently linearly with $\psi_{t,x}$.

The volume of the organs according to their length can be computed from the formulas given in appendix C. The maximal length increase can follow either a linear or a negative exponential function [67]. $Vol_{org,child}$ corresponds to the volume of the child (lateral) organs attached to the node i and which are too small to be explicitly represented in the plant structure. They therefore use the sucrose of parent organ's node they are connected to.

The growth zone of roots are located at the organs' tip [7], while the leaf elongation zone is within the first centimeters at the base of the leaf [68]. For the stem, elongation zone is within the growing phytomer [30]. Therefore:

$$f_{length} = \frac{L_{segingrowingzone}}{L_{growingzone}} \quad (70)$$

After running the phloem module, the water- and carbon-limited growth rate ($r_{CWlim,i}$, $cm^3\ d^{-1}$) for the node $i$ is computed thus:

$$r_{CWlim,i} = \frac{G_{tot,CWlim,i} \times Y}{\rho_s} \quad (71)$$

with $\rho_s$ ($mmol\ sucrose\ cm^{-3}$) the plant tissue sucrose density, $Y$ ($-$) the efficiency of the plant sucrose usage for growth. $G_{tot,i}$ ($mmol\ sucrose\ d^{-1}$) is the actual (carbon- and water-limited) plant sucrose usage rate for growth and growth-related respiration given by the sucrose module (see Eq.(52)). $r_{CWlim,i}$ of the nodes in the growing zone (cell division and cell elongation) will then be used by CPlantBox to simulate the organ growth.
For leaves and stems, this representation creates a small growth-compensation (artificial carbon flow) within the organ's growing zone (first centimeters of the leaf or growing phytomer of the stem). For example: carbon used for growth at the start of the growth section can lead to an elongation at the end of the growth section.
A precise description of the methods used in CPlantBox to elongate or create segments is presented in Schnepf et al. [53].

### F Assumption of local water equilibrium

Thompson and Holbrook [60] presented a mathematical model to justify the assumption of water equilibrium between the xylem and the phloem. Briefly, when their $\hat{R}\hat{F}$ coefficient is above unity, equilibrium can be assumed.
In their model, they looked at a sieve tube with a constant radius and viscosity, where the loading and unloading of sucrose occurred at the boundaries. Those simplification can be applied to our model when focusing on the segment level. Indeed, the (un)loading of sucrose occurs in the nodes at the segments' extremities. The $\hat{R}\hat{F}$ coefficient can be computed thus:

$$\hat{R}\hat{F}_{ij} = \frac{Per_{st}}{A_{cross,st}} \times R\ T_{seg}\ s_{st,.} \times \frac{l_{seg}}{j_{ij}} \times k_{lat,st-x,w} \tag{72}$$

$$s_{st,.} = \begin{cases} s_{st,j} & \text{if} \quad j_{ij} \geq 0 \\ s_{st,i} & \text{else} \end{cases} \tag{73}$$

with $k_{lat,st-x,w}$ the water permeability of the membrane linking the xylem and phloem tissues. In order to have $\hat{R}\hat{F} \geq 1$ we had to adapt our parameter set to have $Fu = 0\ mmol\ ml^{-1}\ d^{-1}$ (no usage of carbon) when $s_{st} < 0.4\ mmol\ ml^{-1}$. This fits with observations for wheat plant where $s_{st}$ was observed to be above this value [60]. For our four simulations, after a burn-in time of 3 $hrs$, we needed to have an area-specific conductance between the xylem and the phloem $k_{lat,st-x,w}$ at least equal to $1.1e-4\ cm\ d^{-1}\ hPa^{-1}$ so that $\hat{R}\hat{F} \geq 1$ for all segments. This value was found to be plausible for wheat plants [6]. This confirms the analysis of Thompson and Holbrook [60]: in their study, they found this simplification to be acceptable for the transport system of most plants, including wheat.

Likewise, in the study of Lacointe and Minchin [33] for $s_{st} > 0.15\ mmol\ cm^{-3}$, decreasing the permeability of the membrane between the xylem and phloem had no effect on the outputs. The assumption of local water equlibrium between xylem and phloem was also tested in experimental studies and found to be acceptable [56, 8, Section 2.2.1.5].

### G Calibration and notations

#### G.1 Experiments for direct calibration

##### G.1.1 Plant growth condition

The wheat phenotype studied in this experiment (UQR15) was obtained from the Hickey lab (University of Queensland, Australia) [48]. 13 PVC columns (diameter = 8 cm, height = 45 cm) were filled with silty loam soil from one testing site of the Research Center Jülich, located in Selhausen (50.8659 N, 6.4471 E). The columns received two seeds each and were afterwards installed in a climate chamber (20°C day, 18°C night, 50% RH, 1000 $\mu mol\ m^{-2}s^{-1}$ of $PAR$ from 6am to 8pm). The mean initial soil gravimetric water content ($\theta_{soil}$) was of 0.40 $cm^3\ cm^{-3}$. Water was added twice a week to keep the $\theta_{soil}$ value between 0.30 $cm^3\ cm^{-3}$ to 0.44 $cm^3\ cm^{-3}$. Fertiliser (7% Nitrate, 19% Ammonium) was added at 6 and 34 days after sowing (stage Z11 and Z25 of the Zadok's growth scale, respectively) in liquid solution supplied at a rate of 3 $kg\ m^{-2}$.

##### G.1.2 Non-destructive measurement

The plant development was evaluated two to three times a week via the following measurements: number of tillers and leaves, plant stage (on the Zadok scale), and length and width of the leaves' blade. The chlorophyll content of the lowest and second highest leaf of each plant tiller was also recorded using a chlorophyll meter (SPAD502, Konika Minolta, Bremen, Germany). The mean $\theta_{soil}$ value for each column was measured by weighting the column on a scale.

##### G.1.3 Soil Water Profiler

In order to measure the transpiration rate, we used the Soil Water Profiler (SWaP) developed by van Dusschoten et al. [65]. At 4, 5, and 6 weeks after sowing (stages Z24, Z25, and Z33, respectively) four of the soil columns were randomly selected. The measurements lasted 36hrs (two day- and one night-periods). The sensor temporal and spatial (vertical) resolutions were 15 min and 1cm, respectively. The day-light intensity alternated between high and low irradiation level every 4hrs. The variation in total $\theta_{soil}$ allowed us to measure the root water uptake rate (assumed equal to the transpiration) at those irradiation levels. The soil surface was covered and the evaporation was thus assumed to be negligible. The same transpiration measurements were done while progressively increasing the irradiation level in order to measure the light saturation point.

##### G.1.4 Destructive measurement

Following the SWaP measurements, we extracted the root systems from the soil. The fresh weight of the roots, stems and leaves was measured. The roots were then stored in 30% Ethanol solutions. The fresh weight of the root, stems and leaf was measured via a weighing scale (precision = 1 mg; Kern 572-30, Kern & Sohn GmbH, Balingen, Germany). The stems and leaves were stored in a freezer at $-20°C$. Afterwards, the leaves were scanned (Canon, 4 Krefeld, Germany) for analysis with ImageJ [52] to measure the leaf blade and sheath surface together with the position and number of the nodes along the stems. The volume of the leaves and stems were obtained by immersing them in a graduated cylinder containing water and measuring the variation of the water level. lateral cuts of the leaf blades (at 5cm to the leaf sheath, on the second leaf of the tiller), stems (at 1cm above the soil, when the stems were present) and roots (at a 9-18cm depth) were done with a cryomicrotome (CM 3050S; Leica Microsystems GmbH, Wetzlar, Germany). The images were analysed with ImageJ to measure the number of vascular bundles and the average shape of the xylem and phloem tissues in each bundle.

#### G.2 Input parameters and other variables and parameters

The tables below present the input parameters and variables of the model. The source used for the parameter estimation can be either [a] direct calibration—direct experimental measurements (see section above), [b] indirect calibration—value set to fit with objective outputs, [c] from the literature (copying the value or recomputing it from the dataset of the publication), in which case the reference is given. The parameters of the CPlantBox core modules are defined by a mean and standard deviation while we only take into account the mean value for the other parameters. The parameters for the organs' shape (the CPlantBox core modules [67]) can be found on the Github repository: https://github.com/Plant-Root-Soil-Interactions-Modelling/CPlantBox/blob/master/modelparameter/structural/plant/Triticum_aestivum_adapted_2023.xml. The shoot shape parameters were obtained from the experiments described above while the root shape parameters (Except for the phloem and xylen tissues) were taken from Bingham and Wu [4].

Table 3: Notation, symbols, values and units - input parameters of the new CPlantBox modules

| Symbol | Description | Values and units | Source | Remarks |
|---|---|---|---|---|
| | | Photosynthesis | | |
| $Chl$ | leaf chlorophyll content | $50.5 \times 10^{-3}$ $\mathrm{mmol^{-1}\,cm^{-2}}$ | direct calibration | |
| $E_{a,c}$ | activation energy for $M_{co2}$ | $59\,430\,\mathrm{mJ\,mmol^{-1}}$ | [64] | |
| $E_{a,j}$ | activation energy $J_{max}$ | $37\,000\,\mathrm{mJ\,mmol^{-1}}$ | [64] | |
| $E_{a,o}$ | activation energy $M_{o2}$ | $36\,000\,\mathrm{mJ\,mmol^{-1}}$ | [64] | |
| $E_{a,rd}$ | activation energy for $R_d$ | $53\,000\,\mathrm{mJ\,mmol^{-1}}$ | [64] | |
| $E_{a,v}$ | activation energy $F_{cmax}$ | $58\,520\,\mathrm{mJ\,mmol^{-1}}$ | [64] | |
| $E_{d,j}$ | deactivation energy $J_{max}$ | $220\,000\,\mathrm{mJ\,mmol^{-1}}$ | [64] | |
| $E_{d,v}$ | deactivation energy $F_{cmax}$ | $220\,000\,\mathrm{mJ\,mmol^{-1}}$ | [64] | |
| $f_{w1r}$ | residual water scarcity factor of the stomatal model | $9.308 \times 10^{-2}$ – | [12] | computed from their dataset |
| $g_0$ | residual stomatal conductance to $CO_2$ | $69\,\mathrm{mmol\,cm^{-2}\,d^{-1}}$ | indirect calibration | from night transpiration |
| $g_{h2o,ox-stomata}$ | conductance of the pathway between the xylem surface and the stomata | $86.4\,\mathrm{mmol\,cm^{-2}\,d^{-1}}$ | indirect calibration | |
| $k_{chl1}$ | fitting parameter for Eqn.(5) | $2\,\mathrm{d^{-1}}$ | indirect calibration | from nitrogen-limited transpiration |
| $k_{chl2}$ | fitting parameter for Eqn.(5) | $9\,\mathrm{mmol\,cm^{-2}\,d^{-1}}$ | indirect calibration | from nitrogen-limited transpiration |
| $k_{fw1}$ | plant sensibility to water stress for stomatal opening | $6 \times 10^{-4}\,\mathrm{hPa}$ | [12] | |
| $k_{g1}$ | fitting parameter for Eqn.(21) | 1.5 – | indirect calibration | from transpiration |
| $k_{g2}$ | $H_2O$ to $CO_2$ ratio of molecular diffusivity in air | 1.6 – | [64] | |
| $k_{jmax}$ | $J^{25}_{max}$ to $F^{25}_{cmax}$ ratio | 1.5 – | indirect calibration | from light saturation point |
| $l_{rootExchange}$ | root exchange (immature) zone | 0 to 0.8 $cm$ from the tip | direct | zone without root hair |
| $M^{25}_{co2}$ | reference Michaelis coefficient for $CO_2$ at 25°C | $302 \times 10^{-6}$ $\mathrm{mmol\,mmol^{-1}}$ | [64] | |
| $M^{25}_{o2}$ | reference Michaelis coefficient for O2 at 25°C | $256 \times 10^{-3}$ $\mathrm{mmol\,mmol^{-1}}$ | [64] | |
| $Mm_{h2o}$ | molar mass of $H_2O$ | $18 \times 10^{-3}\,\mathrm{kg\,mmol^{-1}}$ | - | |
| $o_i$ | substomatal $O_2$ molar fraction | $210 \times 10^{-3}$ $\mathrm{mmol\,mmol^{-1}}$ | [64] | |
| $S$ | entropy term for $J_{max}$ and $F_{cmax}$ | $700\,\mathrm{mJ\,mmol^{-1}\,K^{-1}}$ | [64] | |
| $\alpha$ | fitting parameter for Eqn.(12) | 0.2 – | [64] | |
| $\gamma_0$ | fitting parameter for Eqn.(7) | $28 \times 10^{-6}$ $\mathrm{mmol\,mmol^{-1}}$ | [64] | |
| $\gamma_1$ | fitting parameter for Eqn.(7) | $5.09 \times 10^{-2}\,\mathrm{K^{-1}}$ | [64] | |
| $\gamma_2$ | fitting parameter for Eqn.(7) | $1 \times 10^{-3}\,\mathrm{K^{-2}}$ | [64] | |
| $\psi_{o,symplasm}$ | osmotic water potential in the symplasm | $10\,000\,\mathrm{hPa}$ | [54] | |

| | | | | |
|---|---|---|---|---|
| $\psi_{t,crit,1}$ | critical total water potential of the leaf xylem for stomatal opening | $-2500\,\mathrm{hPa}$ | indirect calibration | |
| $\omega$ | fitting parameter for Eqn.(12) | 0.95 | [64] | |
| | | Organ-specific | | |
| $k_{lat,st,root}$ | root membrane permeability to sucrose | $1 \times 10^{-2}\ \mathrm{cm\,d^{-1}}$ | indirect calibration | to get realistic ratio of exudated sucrose |
| $k_{lat,x,root0}$ | xylem membrane permeability to water for 0-order roots | $6.37 \times 10^{-5}$ $\mathrm{cm\,hPa^{-1}d^{-1}}$ | direct calibration | Based on the concentric membrane model of [7] |
| $k_{lat,x,root1}$ | xylem membrane permeability to water for 1-order roots | $7.9 \times 10^{-5}$ $\mathrm{cm\,hPa^{-1}d^{-1}}$ | direct calibration | |
| $k_{lat,x,root2}$ | xylem membrane permeability to water for 2-order roots | $7.9 \times 10^{-5}$ $\mathrm{cm\,hPa^{-1}d^{-1}}$ | direct calibration | |
| $k_{lat,x,stem}$ | xylem membrane permeability to water for stems | $0\ \mathrm{cm\,hPa^{-1}d^{-1}}$ | - | - |
| $k_{lat,x,leaf}$ | leaf xylem membrane permeability to water | $3.83 \times 10^{-4}$ $\mathrm{cm\,hPa^{-1}d^{-1}}$ | [37] | |
| $r_{st,max,leaf}$ | growth rate used by the phloem module for leaves | $10\,\mathrm{cm\,d^{-1}}$ | indirect calibration | |
| $r_{st,max,root0}$ | growth rate used by the phloem module for 0-order roots | $14.4\,\mathrm{cm\,d^{-1}}$ | indirect calibration | to recreate observed growth once carbon-limited module is used |
| $r_{st,max,root1}$ | growth rate used by the phloem module for 1-order roots | $9\,\mathrm{cm\,d^{-1}}$ | indirect calibration | |
| $r_{st,max,root2}$ | growth rate used by the phloem module for 2-order roots | $8\,\mathrm{cm\,d^{-1}}$ | indirect calibration | |
| $r_{st,max,stem}$ | growth rate used by the phloem module for stems | $1\,\mathrm{cm\,d^{-1}}$ | indirect calibration | |
| $\rho_{s,leaf}$ | sucrose content per volume of leaf fresh tissue | $6.72\,\mathrm{mmol\,cm^{-3}}$ | direct calibration | |
| $\rho_{s,root}$ | sucrose content per volume of root fresh tissue | $6.12\,\mathrm{mmol\,cm^{-3}}$ | direct calibration | |
| $\rho_{s,stem}$ | sucrose content per volume of plant fresh tissue | $7.8\,\mathrm{mmol\,cm^{-3}}$ | direct calibration | |
| | | Other | | |
| $F_{in,max}$ | maximum loading rate into the sieve tube | $0.5\,\mathrm{mmol\,d^{-1}}$ | indirect calibration | |
| $g$ | gravitational acceleration | $980\,\mathrm{hPa cm^2 kg^{-1}}$ | - | |
| $k_{m1}$ | fitting parameters for $R_{m,max}$ | $0.25$ – | indirect | |
| $k_{m2}$ | fitting parameters for $R_{m,max}$ | $2 \times 10^{-5}$ – | indirect calibration | |
| $M_{meso}$ | Michaelis-Menten coefficient for the active loading of sucrose | $0.2\,\mathrm{mmol\,cm^{-3}}$ | indirect calibration | to get realistic sucrose partitioning |
| $M_{out}$ | Michaelis-Menten coefficient for the active unloading of sucrose | $0.1\,\mathrm{mmol\,cm^{-3}}$ | indirect calibration | |
| $Q10$ | temperature coefficient for $R_m$ | 2 – | [3] | |
| $R$ | ideal gas constant | 83.14 $\mathrm{hPa\ cm^3K^{-1}mmol^{-1}}$ | | |
| $s_{st,min}$ | minimum sucrose concentration in the sieve tube below which no loss of carbon occurs | $0.4\,\mathrm{mmol\,cm^{-1}}$ | indirect calibration | |

| | | | |
|---|---|---|---|
| $s_{soil}$ | mean soil soluble carbon concentration in sucrose equivalent content | $1 \times 10^{-4}$ mmol cm$^{-1}$ | direct calibration |
| $T_{ref,Q10}$ | reference temperature for $Q10$ | 293.15 K | [3] |
| $Y$ | plant sucrose use efficiency for growth | 0.9 – | indirect |
| $\beta$ | ratio of axial conductance with sieve plates to that without | 1 – | indirect calibration |
| $\beta_{meso}$ | down-regulation factor for the active loading of sucrose | 0.1 – | indirect calibration |
| $\rho_{h2o}$ | water density | $1 \times 10^{-3}$ kg cm$^{-3}$ | - |
| $\psi_{t,crit,2}$ | critical total water potential of the xylem for growth | $-2000$ hPa | [68] |
| $\psi_{t,max}$ | water potential above which water is not limiting for growth | 0 hPa | [68] |

Table 4: Xylem tubes' shapes

| organ type | tube(s) per vascular bundle | bundle(s) per cross section | resulting $n_i$ | xylem radius, $a_{x,i}$ (cm) | Source |
|---|---|---|---|---|---|
| leaf | 2 | 32 | 64 | 0.0015 | direct calibration |
| leaf | 2 | 32 | 64 | 0.0005 | direct calibration |
| stem | 3 | 52 | 156 | 0.0017 | direct calibration |
| stem | 1 | 52 | 52 | 0.0008 | direct calibration |
| root0 | 4 | 1 | 4 | 0.0015 | direct calibration |
| root1, root2 | 4 | 1 | 4 | 0.00041 | direct calibration |
| root1, root2 | 1 | 1 | 1 | 0.00087 | direct calibration |

Table 5: Sieve tubes' shapes

| organ type | tube(s) per vascular bundle | bundle(s) per cross section | resulting $n_i$ | sieve tube radius, $a_{st,i}$ (cm) | Source |
|---|---|---|---|---|---|
| leaf | 18 | 32 | 576 | 0.00025 | direct calibration |
| stem | 21 | 52 | 1092 | 0.00019 | direct calibration |
| root0 | 33 | 1 | 33 | 0.00039 | direct calibration |
| root1, root2 | 25 | 1 | 25 | 0.00035 | direct calibration |

Table 6: Soil parameters

| Symbol | Description | Values and units | Source |
|---|---|---|---|
| $\theta_{lat,soil}$ | residual water content | 0.08 cm$^3$cm$^{-3}$ | illustrative |
| $\theta_{s,soil}$ | saturated water content | 0.43 cm$^3$cm$^{-3}$ | illustrative |
| $\alpha_{soil}$ | parameter related to the inverse of the air entry suction | 0.04 hPa$^{-1}$ | direct calibration |
| $n_{soil}$ | measure of the pore-size distribution | 1.503 – | illustrative |
| $\kappa_{s,soil}$ | soil conductivity at saturation | 1.6 cm$^2$ hPa$^{-1}$ d$^{-1}$ | illustrative |

Table 7: Notation, symbols and units - variables and other parameters

| Symbol | Description | Units |
|---|---|---|
| | Photosynthesis | |
| $A_g$ | gross carbon assimilation rate per unit of surface | $\mathrm{mmol\,cm^{-2}\,d^{-1}}$ |
| $A_{g,q}$ | daily gross carbon assimilation | $\mathrm{mmol\,d^{-1}}$ |
| $c_i$ | substomatal $CO_2$ molar fraction | $\mathrm{mmol^{-1}\,mmol^{-1}}$ |
| $c_{atm}$ | $CO_2$ molar fraction at the site of measurment | $\mathrm{mmol^{-1}\,mmol^{-1}}$ |
| $D$ | vapor deficit coefficient | $-$ |
| $E_v$ | water vapor flow rate per leaf area | $\mathrm{cm^3\,cm^{-2}\,d^{-1}}$ |
| $ea$ | actual vapor pressure | hPa |
| $es$ | saturated vapor pressure | hPa |
| $f_{w1}$ | water scarcity factor of the stomatal model | $-$ |
| $g_{co2}$ | conductance to $CO_2$ | $\mathrm{mmol\,cm^{-2}\,d^{-1}}$ |
| $g_{ho2}$ | conductance to HO2 | $\mathrm{mmol\,cm^{-2}\,d^{-1}}$ |
| $h$ | relative humidity | $-$ |
| $Je$ | electron transport rate for a set for a set $PAR$ | $\mathrm{mmol\,cm^{-2}\,d^{-1}}$ |
| $Je_{max}$ | maximal electron transport rate | $\mathrm{mmol\,cm^{-2}\,d^{-1}}$ |
| $Je_{max}^{25}$ | maximal electron transport rate at 25°C | $\mathrm{mmol\,cm^{-2}\,d^{-1}}$ |
| $M_{co2}$ | Michaelis coefficient for $CO_2$ | $\mathrm{mmol\,mmol^{-1}}$ |
| $M_{o2}$ | Michaelis coefficient for O2 | $\mathrm{mmol\,mmol^{-1}}$ |
| $P_{atm}$ | atmospheric air pressure | hPa |
| $PAR$ | photosynthetic active radiation | |
| | absorbed by the plant | $\mathrm{mmol\,photons\,cm^{-2}\,d^{-1}}$ |
| $R_d$ | dark respiration rate | $\mathrm{mmol\,cm^{-2}\,d^{-1}}$ |
| $Res$ | resistivity | $\mathrm{d\,cm^2\,cm^{-3}}$ |
| $F_c$ | carboxylation rate | $\mathrm{mmol\,cm^{-2}\,d^{-1}}$ |
| $F_{cmax}$ | maximal carboxylation rate | $\mathrm{mmol\,cm^{-2}\,d^{-1}}$ |
| $F_{cmax}^{25}$ | maximal carboxylation rate at 25°C | $\mathrm{mmol\,cm^{-2}\,d^{-1}}$ |
| $V_j$ | electron transport rate | $\mathrm{mmol\,cm^{-2}\,d^{-1}}$ |
| $\Gamma$ | $CO_2$ compensation point | $\mathrm{mmol\,mmol^{-1}}$ |
| $\Gamma*$ | $CO_2$ compensation point | |
| | in the absence of mitochondrial respiration | $\mathrm{mmol\,mmol^{-1}}$ |
| | Other | |
| $a$ | element's radius or leaf thickness | cm |
| $A_{cross}$ | cross-sectional area | $\mathrm{cm^2}$ |
| $Exud$ | sucrose sink for exudation | $\mathrm{mmol\,d^{-1}}$ |
| $[Exud]$ | sucrose sink for exudation | $\mathrm{mmol\,cm^{-3}\,d^{-1}}$ |
| $F_{in}$, $F_{out}$, $F_{out,MM}$ | source terms for $s_{st}$ loading, total unloading, and unloading with Michaelis-Menten | $\mathrm{mmol\,d^{-1}}$ |
| $[F_{in}]$, $[F_{out}]$, $[F_{out,MM}]$ | source terms for $s_{st}$ loading, total unloading, and unloading with Michaelis-Menten | $\mathrm{mmol\,cm^{-3}\,d^{-1}}$ |
| $G_{tot}$ | sucrose sink for growth | $\mathrm{mmol\,d^{-1}}$ |
| $[G_{tot}]$ | sucrose sink for growth | $\mathrm{mmol\,cm^{-3}\,d^{-1}}$ |
| $j_{lat}$ | solution lateral flux | $\mathrm{cm^1 d^{-1}}$ |
| $J$ | solution volumetric flow rate | $\mathrm{cm^3 d^{-1}}$ |
| $K_{ax}$ | intrinsic axial conductance for the solution | $\mathrm{cm^4 hPa^{-1} d^{-1}}$ |
| $k_{lat}$ | area specific conductance of the membrane | $\mathrm{cm\,hPa^{-1} d^{-1}}$ |
| $L, l$ | total length, arbitrary small length | cm |

| | | |
|---|---|---|
| $Per$ | perimeter | cm |
| $r$ | growth rate | $\text{cm}\,\text{d}^{-1}$ |
| $R_m$ | sucrose sink for maintenance | $\text{mmol}\,\text{d}^{-1}$ |
| $[R_m]$ | sucrose sink for maintenance | $\text{mmol}\,\text{cm}^{-3}\,\text{d}^{-1}$ |
| $s, \bar{s}$ | sucrose concentration, residual between actual and minimum sucrose concentration | $\text{mmol}\,\text{cm}^{-3}$ |
| $s_q$ | sucrose content | mmol |
| $t$ | time | d |
| $Vol$ | volume | $\text{cm}^3$ |
| $T$ | temperature | K |
| $z$ | height, z = 0 at the soil surface, upward-pointing axis | cm |
| $\mu$ | dynamic viscosity | hPa d |
| $\xi$ | local axial coordinate of the plant | cm |
| $\psi$ | water potential | hPa |
| $\Omega$, $\partial\Omega$ | studied domain and its boundaries | − |
| | Subindices | |
| 0 | initial or residual value | |
| $atm$ | atmosphere | |
| $ax$ | axial | |
| $bl$ | leaf boundary layer | |
| $blade$ | leaf blade | |
| $canopy$ | canopy | |
| $Clim$ | carbon limited | |
| $CWlim$ | carbon and water limited | |
| $crit$ | critical | |
| $h$ | hydrostatic (in phloem: turgor) | |
| $g$ | gravitational | |
| $i, j$ | plant node index | |
| $in$ | entering the sieve tube | |
| $l$ | value over the length l | |
| $lat$ | lateral | |
| $lim$ | limited | |
| $m$ | matric potential | |
| $max$ | maximum (potential) value | |
| $meso$ | mesophyll | |
| $o$ | osmotic potential | |
| $ox$ | outer xylem | |
| $org$ | organ | |
| $out$ | outside of plant (value) or leaving the plant (flow) | |
| $p$ | pressure | |
| $sheath$ | leaf sheath | |
| $ref$ | reference | |
| $sheath$ | leaf sheath | |
| $soil$ | soil | |
| $seg$ | plant segment (discretized model) studied plant zone (continuous model) | |
| $st$ | phloem sieve tube | |
| $stomata$ | stomata | |

| | |
|---|---|
| $t$ | total |
| $x$ | xylem |
| $wlim$ | water limited |

### H Numerical solution

In this section, we first give an overview of the main notations used to describe the numerical solutions. We then define the analytical solution for the xylem water flow, leaf outer-xylem water flow (transpiration), and phloem flow for one segment. Finally, we describe the matrix notation and solving scheme. Numerical details of the soil water flow are described in Koch et al. [31].

#### H.1 Mathematical definitions

The branched plant structure is divided into connected line segments with $Ns$ segments and $Nn$ nodes. From the point of view of a mathematical graph, it corresponds to a directed tree (see section 1). According to the graph (or mesh) terminology, a CPlantBox-node is a graph-node (mesh-vertex) and a CPlantBox-segment is a graph-edge ( mesh-cell). For the definition of segment or node variables: We define the segment $ij$ as the segment going from $i$ to $j$.
We have for the segment flows: $J_{ij} = -J_{ji}$ For other segment variables, $X_{ij} = X_{ji}$. Subscripts of brackets or parenthesis ($[...]_{ij}$, $(...)_{ij}$) apply to all the variables within. For the definition of matrices, the subscript $_{seg}$ indicate vectors of size $[Ns, 1]$ containing segment data. $_{node}$ indicate vectors of size $[Ns + 1, 1]$ containing node data. The sign "$\cdot$" indicates a hadamard product, "$\oslash$" a hadamard division and $^T$ indicates a transposition.

The FcVB-stomatal regulation variables (such as $A_g$, $g_{co2,stomata}$, see Section 2.1), the xylem water flux variables ($J_{ax,x}$, $j_{lat,x-soil}$, $j_{lat,x-atm}$, see Section 2.2) and the phloem solution axial flux ($J_{ax,st}$, see Section 2.3) are solved for each segment. The other sucrose-related variables ($s_{st}$, $s_{meso}$, $F_{in}$, $F_{out}$ see Sections 2.3, 2.4) and $\psi_{t,x}$ (see Section 2.2) are solved for each node.

IM is the incidence matrix of the directed graph (of size [Nn, Ns]) and $IM^T$ is its transposition (of size [Ns, Nn]). Thus we have:

$$IM[j,k] = \begin{cases} -1 & \text{if} \quad \text{a segment k has j as begin-node} \\ 1 & \text{if} \quad \text{a segment k has j as end-node} \\ 0 & \text{else} \end{cases} \tag{74}$$

BM is a boolean matrix of size [Ns, Nn]. It gives for each segment the index of the upflow node:

$$BM[k,j] = \begin{cases} 1 & \text{if} \quad IM[j,k] \times J_{w,seg}[k] < 0 \\ 0 & \text{else} \end{cases} \tag{75}$$

with $J_{w,seg}[k]$ the net water flow of the segment $k$ going from the begin-node to the end-node. Therefore, $BM[k,j] = 1$ only if the water flow in the segment $k$ goes from the node $j$ to the other node of the segment $k$.

#### H.2 Analytical solution for one segment

In this following section, we present the definition of the driving equations of Sections 2.2, 2.3 and 2.1, discretized for one plant segment (graph edge) or node (graph vertex).

##### H.2.1 Plant water flow

###### H.2.1.1 Axial flow

The analytical solution to Eqn.(30) presented below is based on the approach of Landsberg and Fowkes [34] and Meunier et al. [39, Appendix B]. However, contrary to them, [a] we define the xylem axial flow according to $\psi_{p,x}$ and not $\psi_{t,x}$ [b] we use an exponential function instead of an hyperbolic function.
When $k_{lat,x} > 0$ and $K_{ax,x} > 0$ we use the analytical continuous solution

$$\psi_{p,x}(l) = \psi_{out} + d_i e^{\tau l} + d_j e^{-\tau l} \quad (76)$$

$$\psi_{out} = \begin{cases} \psi_{m,soil} \text{ if belowground} \\ \psi_{p,ox} \text{ if leaf} \end{cases} \quad (77)$$

with $\psi_{out}$ $(hPa)$ the water potential outside of the plant, uniform over the axial distance $l$. The above equation is solved with

$$\tau = \sqrt{Per_{org} \times k_{lat,x}/K_{ax,x}} \quad (78)$$

where the constants $d_i$, and $d_j$ $(hPa)$ can be calculated from the boundary conditions.
To discretize the equation for each segment $ij$, the Dirichlet top ($l = 0$), and bottom ($l = L$) boundary conditions are given by the begin- and end-nodes of the segment: $\psi_{p,x}(0) = \psi_{p,x,i}$ and $\psi_{p,x}(L) = \psi_{p,x,j}$. They are inserted into the analytic solution and yield the two equations

$$\begin{pmatrix} 1 & 1 \\ e^{\tau L} & e^{-\tau L} \end{pmatrix}_{ij} \begin{pmatrix} d_i \\ d_j \end{pmatrix}_{ij} = \begin{pmatrix} \psi_{p,x,i} - \psi_{out,ij} \\ \psi_{p,x,j} - \psi_{out,ij} \end{pmatrix}. \quad (79)$$

Each root segment is located in one soil voxel defined by a mean $\psi_{m,soil,ij}$ or $\psi_{p,ox,ij}$. Therefore, $\psi_{out,ij}$ is a constant. From these equations we can calculate the constants $d_{i,ij}$, and $d_{j,ij}$ of the segment $ij$ as

$$\begin{pmatrix} d_i \\ d_j \end{pmatrix}_{ij} = \delta_{ij}^{-1} \begin{pmatrix} e^{-\tau L} & -1 \\ -e^{\tau L} & 1 \end{pmatrix}_{ij} \begin{pmatrix} \bar{\psi}_{p,i} \\ \bar{\psi}_{p,j} \end{pmatrix}_{ij} \quad (80)$$

with

$$\delta_{ij} := [e^{-\tau L} - e^{\tau L}]_{ij}, \quad (81)$$

and

$$\bar{\psi}_{p,i,ij} = \psi_{p,x,i} - \psi_{out,ij} \quad (82)$$

$$\bar{\psi}_{p,j,ij} = \psi_{p,x,j} - \psi_{out,ij} \quad (83)$$

Thus, $\bar{\psi}_{p,i,ij}$ and $\bar{\psi}_{p,j,ij}$ are constants.
Putting the constants $d_{i,ij}$ and $d_{j,ij}$ into the solution for $\psi_{p,x}$ (Eqn.(76)) we can write down the exact axial flow in the segment $ij$, as [39, B2]:

$$J_{ax,x,ij}(l) = -[K_{ax,x} \left(d_i \tau e^{\tau l} - d_j \tau e^{-\tau l} + d\psi_g(l)\right)]_{ij} \quad (84)$$

with $d\psi_g(l)$ the gradient of gravitational water potential between the nodes $i$ and the point at $l$ $cm$ from $i$. Replacing the constants $d_{i,ij}$ and $d_{j,ij}$ with their definition yields the explicit axial flow equation:

$$\begin{aligned} J_{ax,x,ij}(l) = &- K_{ax,x,ij}[\delta^{-1}\tau[(e^{-\tau L}\bar{\psi}_{p,x,i} - \bar{\psi}_{p,x,j})e^{\tau l} \\ &- (-e^{\tau L}\bar{\psi}_{p,x,i} + \bar{\psi}_{p,x,j})e^{-\tau l}] + d\psi_g(l)]_{ij} \end{aligned} \quad (85)$$

Evaluation of the axial flow at $l = 0$ (node $i$) yields

$$J_{ax,x,ij}(0) \quad = \quad -K_{ax,x,ij}\left(\delta^{-1}\tau[(e^{-\tau L} + e^{\tau L})\bar{\psi}_{p,i} - 2\bar{\psi}_{p,j}] + d\psi_g(0)\right)_{ij}. \tag{86}$$

For conservation of mass in each node $i$, the summed exact axial flows have to equal zero,

$$\sum_{j \in N(i)} J_{ax,x,ij}(0) = 0 \tag{87}$$

where N(i) are the indices of the segments starting from node index $i$ to $j$. We can solve this system of equations (see appendix H.3.1) and obtain thus $\psi_{p,x}$ and $J_{ax,x}$ for each plant node and segment, respectively.

$\psi_{p,x}$ and $J_{ax,x}$ are then used by the photosynthesis and phloem module (Eqns.(19) and (32)) and by the core CPlantBox-module to compute the water-limited growth rate (Eqn.(68)).

###### H.2.1.2 Lateral flow

Once $\psi_{p,x}$ is known, the exact lateral water flow ($J_{lat,x-out}$, in $cm^3 d^{-1}$) for a segment $ij$ with length $L_{ij}$ and lateral conductivity $k_{lat,x,ij}$ can be computed with the following equations:

$$J_{lat,x-out,ij} \quad = \quad -[Per_{org}\ k_{lat,x}]_{ij} \int_0^{L_{ij}} \psi_{out,ij} - \psi_{p,x} dl \tag{88}$$

$$J_{lat,x-out,ij} \quad = \quad [Per_{org}\ k_{lat,x}\left(\frac{d_i}{\tau}(e^{\tau L} - 1) + \frac{d_i}{\tau}(1 - e^{-\tau L})\right)]_{ij} \tag{89}$$

Inserting the definition of $d_{i,ij}$ and $d_{i,ij}$ into the exact lateral flow yields

$$\begin{aligned} J_{lat,x-out,ij} = & [Per_{org}\ k_{lat,x}\frac{1}{\delta\tau} \\ & \times \left(\bar{\psi}_{p,i}(1 - e^{-\tau L} - e^{\tau L} + 1) + \bar{\psi}_{p,j}(-e^{\tau L} + 1 + 1 - e^{-\tau L})\right)]_{ij} \end{aligned} \tag{90}$$

and

$$J_{lat,x-out,ij} \quad = \quad [Per_{org}\ k_{lat,x}\frac{1}{\delta\tau}(\bar{\psi}_{p,i} + \bar{\psi}_{p,j})(2 - e^{-\tau L} - e^{\tau L})]_{ij} \tag{91}$$

The $J_{lat,x-out}$ values from root segments can then be used as source term in the soil module (see Section 3). The $J_{lat,x-out}$ values from leaf segments corresponds to the transpiration rate (see Section 2.1.1).

##### H.2.2 Stomatal regulation, photosynthesis, and transpiration

Once $\psi_{p,x}$ and $j_{lat,x-atm}$ are known, we can compute $\psi_{p,ox}$ ($hPa$), the water potential on the surface of the leaf vascular bundle-sheath, by implementing Eqns.(90), and solving for $\psi_{out}$ (here, equivalent to $\psi_{p,ox}$):

$$\psi_{p,ox,ij} \quad = \quad \left(\left[\frac{j_{lat,x-atm} \times L \times Per}{k_{lat,x}\frac{1}{\tau\delta}(2 - e^{-\tau L} - e^{\tau L})}\right]_{ij} + (\psi_{h,x,i} + \psi_{h,x,j})\right) \times 0.5 \tag{92}$$

$j_{lat,x-atm}$ is obtained from Eqn.(3). $\psi_{p,ox}$ can then used in the xylem flow module (see Eqn.(29)) at the next computation loop.

##### H.2.3 Sucrose balance

As explained above, $s_{st}$, $s_{meso}$, $F_{in}$, $F_{out}$ are node variables. When segment-defined variables or parameters are needed to compute them (such as $Vol_{st}$, $A_g$, $k_{lat,st}$), we allocate to the node $i$ the variable from the segment .i which has $i$ as end node: $X_{i,node} = X_{.i,seg}$.

We set $s_{Y,q}$ ($mmol\ sucrose$) the sucrose content in the compartment $_Y$, either the mesophyll or the sieve tube.
$\frac{ds_{q,j}}{dt}$ ($mmol\ sucrose\ d^{-1}$), the variation of sucrose content in the node j, is defined thus:

$$\frac{ds_{st,q,j}}{dt} = J_{ax,st,q,j.} + (F_{in,j} - F_{out,j}) \tag{93}$$

$$J_{ax,st,q,j} = \sum_{i}^{Nn} (s_{ij} \times J_{ax,st,h2o,ij}) \tag{94}$$

$$J_{ax,st,h2o,ij} = \frac{K_{ax,st,ij}}{L_{ij}} \times (RT \times (s_j - s_i) + (\psi_{xj} - \psi_{xi})) \tag{95}$$

$$s_{ij} = \begin{cases} s_i & \text{, if} \quad J_{w,ij} < 0 \\ s_j & \text{, if} \quad J_{w,ij} \geq 0 \end{cases}, \tag{96}$$

$$\frac{ds_{meso,q,j}}{dt} = (F_{Ag,i} - F_{in,i}) \times Vol_{meso,i}. \tag{97}$$

$\psi_{xj}$ and $\psi_{xi}$ are obtained thanks to the xylem flow computation (see Sections 2.2 and H.2.1). $F_{Ag,i}$ corresponds to the sucrose assimilation rate computed according to Eqns.(16),(40). $F_{in,i}$ and $F_{out,i}$ are computed according to the equations in Eqns.(42) and (50) respectively. $J_{ax,st,q,j.}$ (resp. $J_{ax,st,h2o,ij}$) corresponds to the axial flow in the sieve tube of sucrose content (resp. water) at the node i.

#### H.3 Matrix form and solving scheme

In the following section, we present the discretized system of equations and the solving schemes to obtain $\psi_{p,x}$, $\frac{ds_{st,q,j}}{dt}$ and $\frac{ds_{meso,q,j}}{dt}$.

##### H.3.1 Xylem water flow

From Eqns.(86-87) we can construct a system of equations, which can be solved for $\psi_{p,x}$:

$$\psi_{h,x,node} = C^{-1} \times b_{node} \tag{98}$$

With $C$ and $b_{node}$ matrices of size $[Nn, Nn]$ and $[Nn, 1]$ respectively. They are filled with zeros except at the following locations:

$$C[i, i] = C[i, i] + c_{i,ij} \tag{99}$$

$$C[i, j] = 2[K_{ax,x}\delta^{-1}\tau]_{ij} \tag{100}$$

$$b_{node}[i, 1] = b_{node}[i, 1] + [K_{ax,x} \times d\psi_g]_{ij} + (c_{i,ij} + C[i, j])\psi_{out,ij} \tag{101}$$

$$c_{i,ij} = [K_{ax,x}\delta^{-1}\tau(e^{-\tau L} + e^{\tau L})]_{ij} \tag{102}$$

for each segment $ij$ going from node $i$ to node $j$.

This system of equation is solved for the steady state solution of $\psi_{p,x}$ with the sparse linear solver (SParseLU) of the C++ Eigen library [23]. The $\psi_{p,x}$ are then used to compute the variables from the FcVB-stomatal regulations module. The fixed point iteration between the computation continues until convergence.

##### H.3.2 Sucrose balance

The equations from Section H.2.3 can be set in matricial form:

$$\left[\frac{ds_{st,q}}{dt}\right]_{node} = IM \times (J_{w,seg} \cdot s_{st,seg}) + (F_{in,node} - F_{out,node}) \cdot Vol_{st,node} \quad (103)$$

$$s_{st,seg} = BM \times s_{st,node} \quad (104)$$

$$s_{st,node} = s_{st,q,node} \oslash Vol_{st,node} \quad (105)$$

$$J_{w,seg} = Co_{ax,st,seg}(IM^T \times RTs_{st,node} + IM^T \times \psi_{x,node}) \quad (106)$$

$$\left[\frac{ds_{meso,q}}{dt}\right]_{node} = (F_{Ag,node} - F_{in,node}) \cdot Vol_{st,node} \quad (107)$$

with $Co_{ax,st,seg}$ the vector defining the conductance ($cm^3hPa^{-1}d^{-1}$) per sieve tube segment.

The system of equations is solved numerically using an implicit method: the value of $s^{m,n}_{q,node}$ (the vector containing the estimation $m$ at time step $n$ of $s_q$ per node) is used to compute $\left[\frac{ds^{m,n}_q}{t_n - t_{n-1}}\right]_{node}$. This value is sent to the implicit Sundials-CVODE solver [25] which then yields $s^{m+1,n}_{q,node}$. This loops continues until convergence is reached. The solver then moves on the the next time step.

### I Sensitivity analysis

The program contains a high number of plant parameters which cannot be measured directly and which are highly correlated between them with regards to the outputs. Therefore, to have a better idea of which parameters need to be (re)calibrated in priority, a first sensitivity analysis was run for [a] the coupled water flow and FcVB-stomatal regulations modules (water modules), [b] the phloem flow and carbon usage modules (carbon modules). A sensitivity analysis of the root system parameters can be found in the work of Morandage et al. [40] .

As the computation time of the full coupled simulation is too long to run a sensitivity analysis, we evaluated the effect of the water (resp. carbon) modules' parameters for one time step of 3 $hrs$. We moreover focused on a selected number of outputs variables of interest, namely [a] the total transpiration and gross assimilation rates for the water modules and [b] carbon usage rate for growth ($G_{tot,CWlim}$) for the carbon modules.

This sensitivity analysis is limited to a short time span and evaluates each module separately. Nonetheless, water (resp. phloem) parameters which have a strong influence on the transpiration and assimilation rates (resp. growth rate) will also influence strongly the other modules on the long term. For this evaluation, we did not include plant parameters which could be considered as physical constants, like $E_a$ and $E_d$, respectively the activation and deactivation energy parameters used in the photosynthesis model.

The sample range of each evaluated parameter was set to [$\frac{1}{2}$,2] of their value used in this study, except for the values of $\alpha$ and $\omega$. For these two variables, ranges were set to make sure that a solution for Eqn. (12) could be found. We used $\Delta\psi_{o,symplast}$ as parameter(which corresponds to $\psi_{o,symplast} - \psi_{t,crit2}$) instead of $\psi_{o,symplast}$ to avoid ever having $|\psi_{o,symplast}| > |\psi_{t,crit2}|$ which would have led to computation errors. Moreover, a first analysis showed that the environmental conditions and the plant's development stage influenced the sensitivity of the outputs to the parameters. Therefore, the analysis was run for each plant age at the beginning of the simulations (11 or 18 days) and each weather type (*drier&warmer*, *wetter&colder*) using the environmental variables for noon.

The analysis was done with the SALib python module [26]. The number of samples was set according to the trade-off between precision gain and computation speed loss (see table 8). Therefore, the simulations using slower modules were run with a lower number of parameter samples.

Table 8: Number of samples per parameter for each simulation for the Sobol sensitivity analysis

| | water modules | | carbon modules | |
|---|---|---|---|---|
| scenario \ plant age | 11d | 18d | 11d | 18d |
| wetter&colder | $2^{11}$ | $2^{11}$ | $2^{8}$ | $2^{8}$ |
| drier&warmer | $2^{11}$ | $2^{11}$ | $2^{8}$ | $2^{8}$ |

Figure 13 presents the outputs of the analysis. The variation of the sensitivity index according to the plant and environment's characteristics indicate that such analysis could be necessary for each model setup. The difference in the width of the confidence intervals is in part linked to the different number of samples used for each analysis (see Table 8).

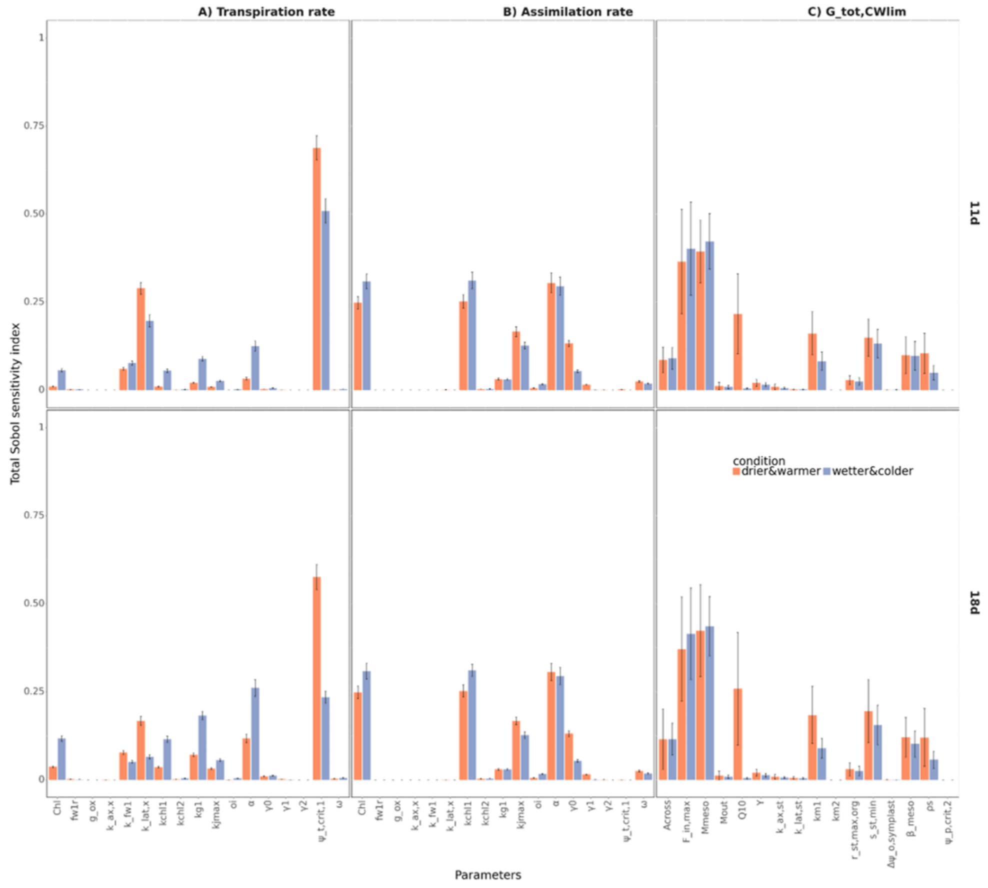


Figure 13: **Sensitivity analysis** of the plant's (A) transpiration and (B) gross assimilation rate to the water modules' parameter and (C) carbon usage rate for growth ($G_{tot,CWlim}$) to the carbon modules' parameters for 11d and 18d plants under the two weather scenarios presented in the study. The sensitivity index includes both first order effects and interactions with the other parameters. The error bars show the limits of the confidence interval (confidence level of 95%).

For the water modules, we can observe that $\psi_{t,crit,1}$ and $k_{fw1}$ are especially important. These parameters define the decrease in leaf lateral conductivity because of low leaf water potential. For the assimilation rate, $Chl$ (the leaf chlorophyll content) and $k_{chl1}$ (a parameter linking the chlorophyll content to the potential assimilation rate), have a strong influence on the output of the simulation, because the affect the carboxylation rate ($V_c$). Similarly, $\alpha$ and $k_{jmax}$ affect the photon-limited assimilation rate $V_j$.

For the carbon modules, we can observe for instance that $G_{tot,CWlim}$ is especially sensible to the parameters defining the flow of sucrose from the mesophyll to the sieve tubes: $F_{in,max}$, $M_{meso}$ and $\beta_{meso}$. This defines how quickly the assimilated sucrose flows in the sieve tube and has a strong influence on the plant's capacity to buffer changes in assimilation rate. The temperature coefficient $Q10$ represent the effect of temperature on $R_m$. For the *warmer&drier*, the simulated temperature at noon is further away from the reference temperature than for *wetter&colder*,

leading to a stronger effect of $Q10$.
Surprisingly, $\psi_{p,crit,2}$ which defines the water limitation on growth, was found to have no influence on the outputs. It is likely caused by the fact that, for the simulated period, sucrose was more limiting than water for growth. Running a longer simulation for the sensitivity analysis should give a better evaluation of the influence of $\psi_{p,crit,2}$ on $G_{tot,CWlim}$.

### J Other simulation Results

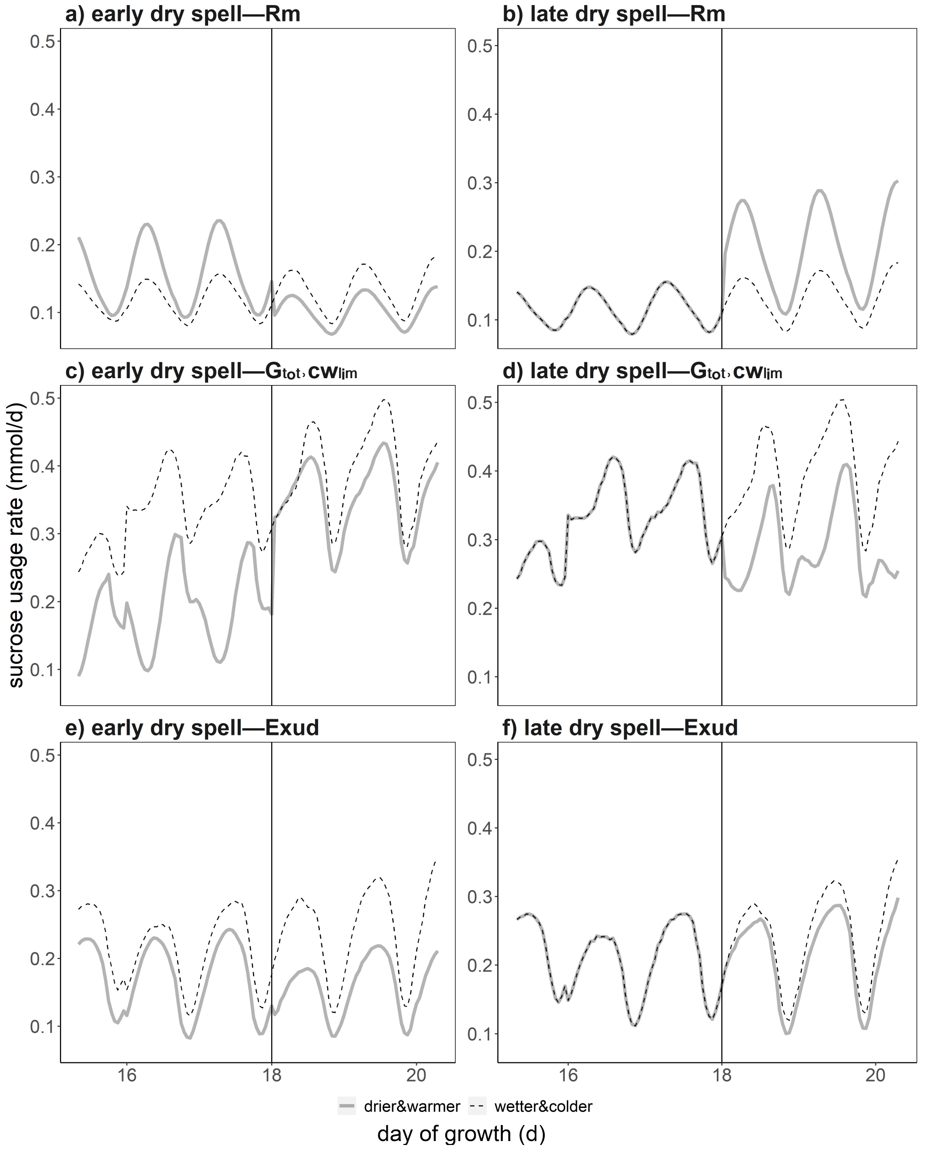


Figure 14: **Partitioning of the carbon loss between the three sinks according to time** between day 15 and 20: maintenance respiration ($Rm$), growth and growth respiration ($G_{tot,CWlim}$), root exudation ($Exud$). The black vertical line defines end (resp. start) of the early (resp. late) dry spell. We compare wetter and colder dry spells (thin dotted lines) against drier and warmer dry spells (thick lines).
